# GPCR-mediated Activation of PLCγ2 Initiates Dysregulated Recruitment of Neutrophils in Cold-induced Urticaria in PLAID Patients

**DOI:** 10.1101/2020.12.22.424011

**Authors:** Xuehua Xu, Xi Wen, Smit Bhimani, Amer Moosa, Dustin Parsons, Tian Jin

**Author notes:** Correspondence to Xuehua Xu.

## Abstract

The current dogma is that chemoattractants G protein coupled receptors (GPCRs) activate β phospholipase C (PLCβ) while receptor tyrosine kinases (RTKs) activate γ phospholipase C (PLCγ). Here, we show that chemoattractant/GPCR-mediated membrane recruitment of PLCγ2 constitutes GPCR-mediated phospholipase C (PLC) signaling and is essential for neutrophil polarization and migration during GPCR-mediated chemotaxis. In response to a chemoattractant stimulation, cells lacking PLCγ2 (*plcg2^kd^*) displayed altered dynamics of diacylglycerol (DAG) production and calcium response; increased Ras/PI3K/Akt activation; elevated GSK3 phosphorylation and cofilin activation; impaired dynamics of actin polymerization; and consequently defects in cell polarization and migration during chemotaxis. At low temperature, neutrophils expressing the gain-of-function mutant of PLCγ2 (Δ686) displayed better chemotaxis than the cells expressing wild-type PLCγ2. The study provides a molecular mechanism for the dysregulated recruitment and activation of neutrophils in cold-induced urticaria in PLCγ2-associated antibody deficiency and immune dysregulation (PLAID) patients bearing gain-of-function mutations of PLCγ2.

## INTRODUCTION

Neutrophil chemotaxis plays critical roles in innate immune responses and forms first-line host defense against infections (Liew and Kubes, 2019; Nathan, 2006). Inappropriate recruitment and dysregulated activation of neutrophils contribute to tissue damage and cause autoimmune and inflammatory diseases (Kolaczkowska and Kubes, 2013; Nourshargh and Alon, 2014). Neutrophils use G protein-coupled receptors (GPCRs) to sense chemoattractants and chase bacteria, while they use phagocytic receptors coupled with tyrosine kinases to engulf and kill bacteria. The current dogma is that chemoattractants/GPCRs activate β phospholipase C (PLCβ) and receptor tyrosine kinases (RTKs) activate γ phospholipase C (PLCγ). In neutrophils, the engagement of chemoattractants with their GPCRs activates β2 and β3 phospholipase C (PLCβ2/β3) through heterotrimeric G proteins (Camps et al., 1992; Jiang et al., 1994; Li et al., 2000; Park et al., 1993). Activated PLCβ2/β3 catalyze the hydrolysis of phosphatidylinositol 4,5 bisphosphate (PI(4,5)P2, PIP2) to yield inositol 1,4,5 trisphosphate (Ins(1,4,5)P3, IP3) and diacylglycerol (DAG), which are important secondary cellular messengers to induce calcium response and activate the effectors of PLC, such as protein kinase C (PKC) in neutrophils. Murine neutrophils with double deficiency of PLCβ2/3 (*plcb2^-/-^ plcb3^-/-^* display decreased PLC signaling, such as reduced IP3 production, PKC activation, and superoxide release, indicating an essential role of PLCβ2/3 in bacterial killing (Li et al., 2000). *plcb2^-/-^ plcb3^-/-^* murine neutrophils chemotax normally toward fMLP and MIP-1α gradients, leading to the assumption that PLCβ and PLC signaling might not be required for GPCR-induced neutrophil chemotaxis (Li et al., 2000). Recently, we and others demonstrated the essential roles of PLCβ and PLC signaling in cofilin/SSHs regulation for neutrophil polarization and chemotaxis (Tang et al., 2011; Xu et al., 2015b). Mammalian neutrophils highly express PLCγ2, in addition to PLCβ2/β3 (Suh et al., 2008). PLCγ2 plays critical roles in integrin- and Fc receptor-mediated neutrophil functions, and its activation is through Syk-mediated phosphorylation (Jakus et al., 2009), consistent with the traditional belief that GPCRs activate β phospholipase C (PLCβ) and receptor tyrosine kinases (RTKs) activate γ phospholipase C (PLCγ) (Suh et al., 2008). PLCγ2-associated antibody deficiency and immune dysregulation (PLAID) and autoinflammation and PLCγ2-associated antibody deficiency and immune dysregulation (APLAID) patients who bear gain-of-function mutations in PLCγ2 and display excessive neutrophil infiltration, a process closely related to the chemotaxis, in cold-induced urticaria (Aderibigbe et al., 2015; Ombrello et al., 2012; Zhou et al., 2012). Inconsistent with the above, the neutrophils purified from PLAID and APLAID patients displayed impaired chemotaxis (Aderibigbe et al., 2015). We found that chemoattractants induce robust membrane translocation of PLCγ2 in human neutrophils (Xu et al., 2015b), a well-characterized mechanism for PLCγ activation (Falasca et al., 1998; Nishida et al., 2003). Chemotaxing human neutrophils actively recruit PLCγ2 to their leading fronts (Xu et al., 2016), implicating a potential function of PLCγ2 in the remodeling of cytoskeletons at the leading front during neutrophil chemotaxis. However, the molecular mechanism of PLCγ2 function in GPCR-mediated PLC signaling and neutrophil chemotaxis and its role in dysregulated recruitment and activation of neutrophils in PLAID patients remain unknown.

In the present study, we show that membrane recruitment and subsequent activation of PLCγ2 during neutrophil chemotaxis requires chemoattractant GPCR-mediated signaling events, constitutes the essential PLC signaling for polarization and migration during chemotaxis. In response to chemoattractant stimulation, cells lacking PLCγ2 (*plcg2^kd^*) display a series of dysregulated GPCR-mediated signaling events and defective chemotaxis. More importantly, we found that, at low temperature, cells expressing gain-of-function mutant of PLCγ2 displayed an improved chemotaxis capability than the cells expressing either empty vector or wild-type PLCγ2. Our work sheds new light on the dogmatic understanding on how different classes of receptors activate PLC isoforms. More importantly, this work provides a molecular mechanism for the initiation of the dysregulated recruitment and activation of neutrophils during cold-induced urticaria in the PLAID patients bearing the PLCγ2 mutation.

## RESULTS

### Molecular mechanism of GPCR-mediated membrane recruitment and activation of PLCγ2

Neutrophils highly express three PLC isoforms: PLCβ2, β3, and γ2 (Fig. S1A) (Suh et al., 2008). Binding of chemoattractants to their GPCRs induces activation/dissociation of heterotrimeric G protein and the free Gα-GTP and Gβγ directly activate PLCβ isoforms (Camps et al., 1992; Jiang et al., 1994; Li et al., 2000; Park et al., 1993). We and others have shown the essential role of PLC signaling in neutrophil chemotaxis (Tang et al., 2011; Xu et al., 2016). However, one previous study showed that murine neutrophils deficient in both *plcb2* and *plcb3* chemotax normally toward fMLP and MIP-1α gradients (Li et al., 2000), indicating the potential role of other PLC isoforms, such as PLCγ2, in neutrophil chemotaxis (Li et al., 2000). Consistent with the study, we found that various chemoattractants induce robust translocation of PLCγ2 to the plasma membrane (PM) (Xu et al., 2015a), an efficient way of activating PLCγ (Falasca et al., 1998; Koss et al., 2014). In the gradients of diverse chemoattractants, PLCγ2 is also actively recruited to the leading front of chemotaxing cells (Xu et al., 2015a). Consistent with previous reports, we detected a similar expression pattern of PLC isoforms in human neutrophil-like (HL60) cells to that in primary mammalian neutrophils (Fig. S1) and identified it as a suitable cell line in which to study the function of PLCγ2 in mammalian neutrophils (Ramirez et al., 2017; Rincon et al., 2018). To understand the molecular mechanism of chemoattractant-mediated PLCγ2 membrane targeting, we investigated the domain requirement for its membrane translocation. PLCγ2 has multiple domains that may contribute to its translocation to the PM. Therefore, we generated GFP-tagged wild-type (WT), lipase dead (LD), and truncated mutants (ΔPH and ΔC2) of PLCγ2 (green) (Fig. S2). To visualize the PM, we used several PM markers in the current study (Fig. S3). We expressed the GFP-tagged PLCγ2 or its mutants (green) along with a PM marker (CAAX-mCherry, red) and monitored their PM translocation in the cells in response to fMLP stimulation (Fig. 1A). We found that WT and mutants of PLCγ2 localized at the protruding sites of resting HL60 cells. In response to a uniformly applied fMLP stimulation, WT PLCγ2 transiently translocated to and colocalized with a PM marker on the entire PM and then continuously accumulated on the protruding sites of the cells (Video S1). In contrast to a robust membrane translocation of WT, all three mutants displayed decreased membrane translocation with different dynamics: LD displayed a noticeably decreased membrane translocation with a continuous translocation to the protruding sites of cells (Video S2); ΔPH displayed a clear initial membrane translocation but a reduced accumulation on protruding sites of the cells (Video S3); and ΔC2 displayed a significant decrease in both the initial and the second membrane translocation (Video S4). Quantitative measurement of the membrane translocation of WT and mutants of PLCγ2 confirmed biphasic membrane translocation of WT and a reduced membrane translocation of the mutants with altered dynamics (Fig. 1B). This result is consistent with previous reports that both PH-PIP_3_ (product of chemoattractant/GPCR activation) and C2/Ca^2+^ binding play a role in the membrane recruitment of PLCγ2 (Falasca et al., 1998; Nishida et al., 2003; Xu et al., 2015b). It has been shown that Integrin and Fc receptors activate PLCγ2 through Src/syk phosphorylation of its tyrosine residue Y759 corresponding to PLCγ1(Y783) in neutrophils (Jakus et al., 2009; Koss et al., 2014). We further examined whether chemoattractants also induce the phosphorylation of PLCγ2 (Fig. 1C). We detected the phosphorylation of PLCγ2 (Y759) in IgG-treated cells, but not in cells stimulated with either chemoattractant fMLP or SDF1α, although all three treatments induced Erk phosphorylation in the cells. This result indicates that chemoattractant stimulation does not trigger tyrosine phosphorylation-mediated activation of PLCγ2. Thus, the above results indicate that chemoattractant-mediated membrane targeting and activation of PLCγ2 requires chemoattractant-induced PIP_3_ production and [Ca^2+^] increase.

**Figure 1.**
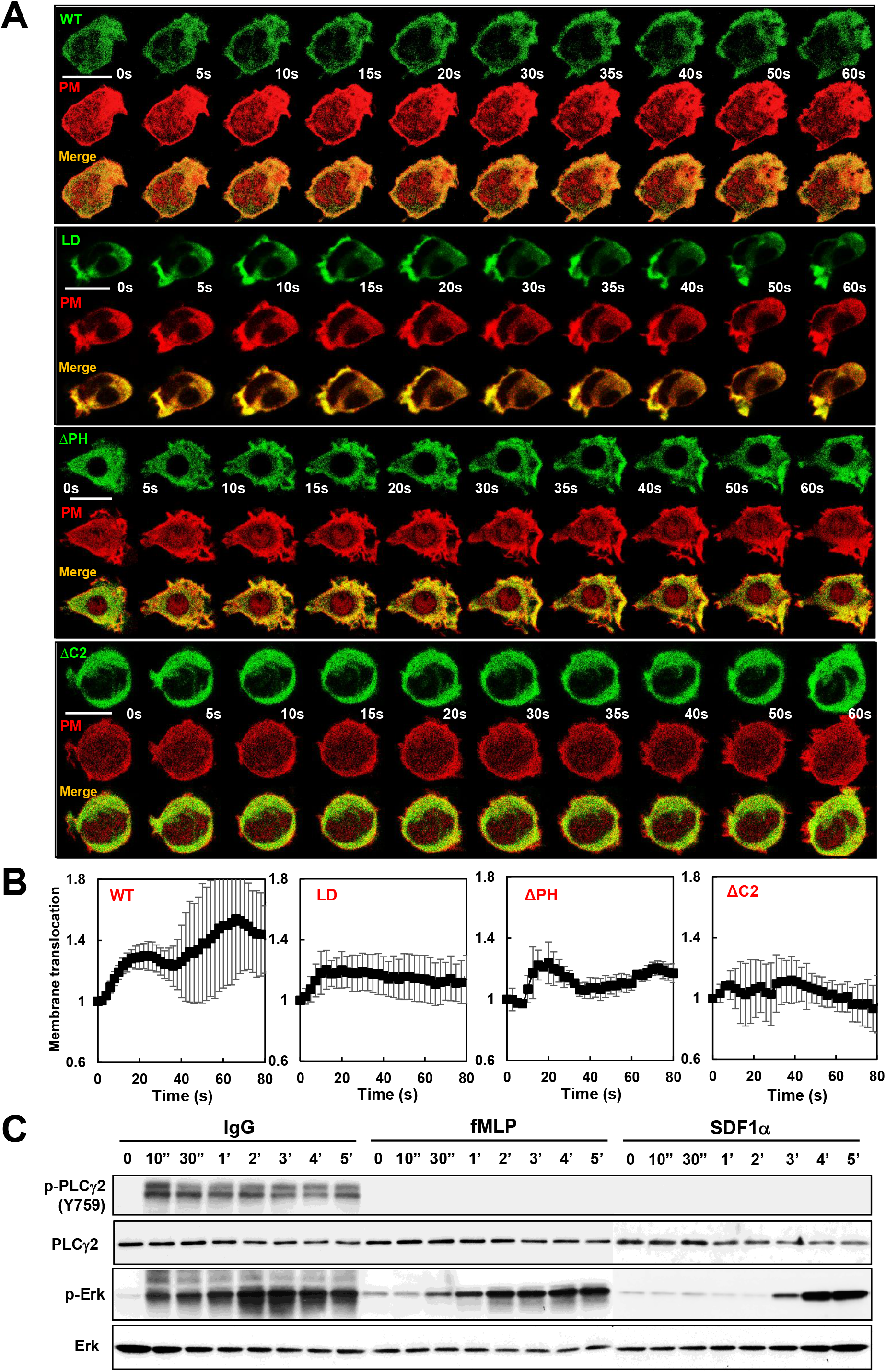
Chemoattractants induces membrane targeting of PLCγ2. (A) Montage shows fMLP-induced membrane translocation of the wild-type (WT) or the mutants of PLCγ2 in response to uniformly applied fMLP stimulation. For complete sets of responses see Video S1-S4. Cells expressing a PM marker (CAAX-mCherry, red) and GFP-tagged WT (Video S1) or mutants of PLCγ2 (green), including lipase dead (LD, Video S2) and truncation mutants lacking either PH (ΔPH, Video S3) or C2 (ΔC2, Video S4) domain, were imaged in time-lapse; 100 nM fMLP was applied to the cells after 0 s. (B) Quantification of the membrane translocation of PLCγ2 WT or mutants in response to fMLP stimulation. Mean ± SD is shown; n= 6, 5, 5, or 5 for WT, LD, ΔPH, and ΔC2 of PLCγ2, respectively. (C) Chemoattractant stimulation does not induce phosphorylation of PLCγ2. At time 0 s, cells were stimulated with 1 μM fMLP, or 1 μg/ml SDF1a, or 1 μg/ml IgG. Aliquots of the cells at indicated time points were lysed and subjected to western blot detection of phosphorylated and total PLCγ2 using their specific antibodies. p-Erk1/2 or total Erk1 was also detected to determine that cells were responsive to chemoattractant stimuli. The same set western blot result detected with the indicated antibodies, including the set of cells stimulated with IgG and fMLP and the set stimulated with IL8, were derived from two membranes and performed with the exact same western-blotting procedure.

### PLCγ2 plays an essential role in the GPCR-mediated PLC activation

To understand how PLCγ2 contributes to GPCR-mediated PLC signaling, we infected HL60 cells with lentiviral particles encoding non-specific or *plcg2* specific shRNA and generated a control (CTL) and a *plcg2^kd^* cell line, in which the expression of *plcg2* was stably knocked down (Fig. 2A). We confirmed that *plcg2^kd^* cells expressed significantly reduced level of PLCγ2 without affecting the expression of chemoattractant receptors, such as FPR1 receptors (Fig. 2B). PLC hydrolyzes PIP2 and generates IP3 and DAG. To examine how PLCγ2 contributes to GPCR-mediated PLC signaling, we first examined DAG production in both CTL and *plcg2^kd^* cells in response to fMLP stimulation by monitoring the membrane translocation of a DAG biosensor DBD-YFP (DAG-binding domain of protein kinase CβII, green) and a PM marker (CAAX-mCherry, red) using fluorescence microscopy (Fig. 2C) (Gallegos et al., 2006; Violin et al., 2003). In response to uniformly applied fMLP stimulation, in CTL cells, DBD-YFP transiently translocated to and colocalized with the PM marker on the entire PM and then continuously localized at the protrusion sites of the migrating cells (Fig. 2C, upper panel, and Video S5). This indicates that uniform chemoattractant stimulation induces a biphasic PLC activation, which is consistent with the biphasic dynamics of PLCγ2 membrane targeting. In *plcg2^kd^* cells, uniform fMLP stimulation also triggered initial DBD-YFP translocation and then returned to the cytoplasm with a reduced secondary accumulation of DBD-YFP in the protruding sites of cells (Fig. 2C, middle panel, and Video S6). We then expressed PLCγ2-CFP (blue) in *plcg2^kd^* cells (*plcg2^kd/OE^*) and monitored DAG production (DBD-YFP, green) along with PM marker (red) in *plcg2^kd/OE^* cells upon uniform fMLP stimulation (Fig. 2C, lower panel, and Video S7). We found that *plcg2^kd/OE^* cells displayed biphasic dynamics of DBD membrane translocation. More importantly, *plcg2^kd/OE^* displayed repolarization and migration behavior similar to that of CTL cells. Quantitative measurement of DBD-YFP membrane translocation in multiple cells showed a typical biphasic DAG production in CTL and *plcg2*^*kd*/OE^ cells, but a single-phase, transient DAG production in *plcg2^kd^* cells (Fig. 2D). The above result indicates that PLCγ2 plays a major role in the second-phase activation of PLC signaling, during which cells polarized and then start to migrate.

**Figure 2.**
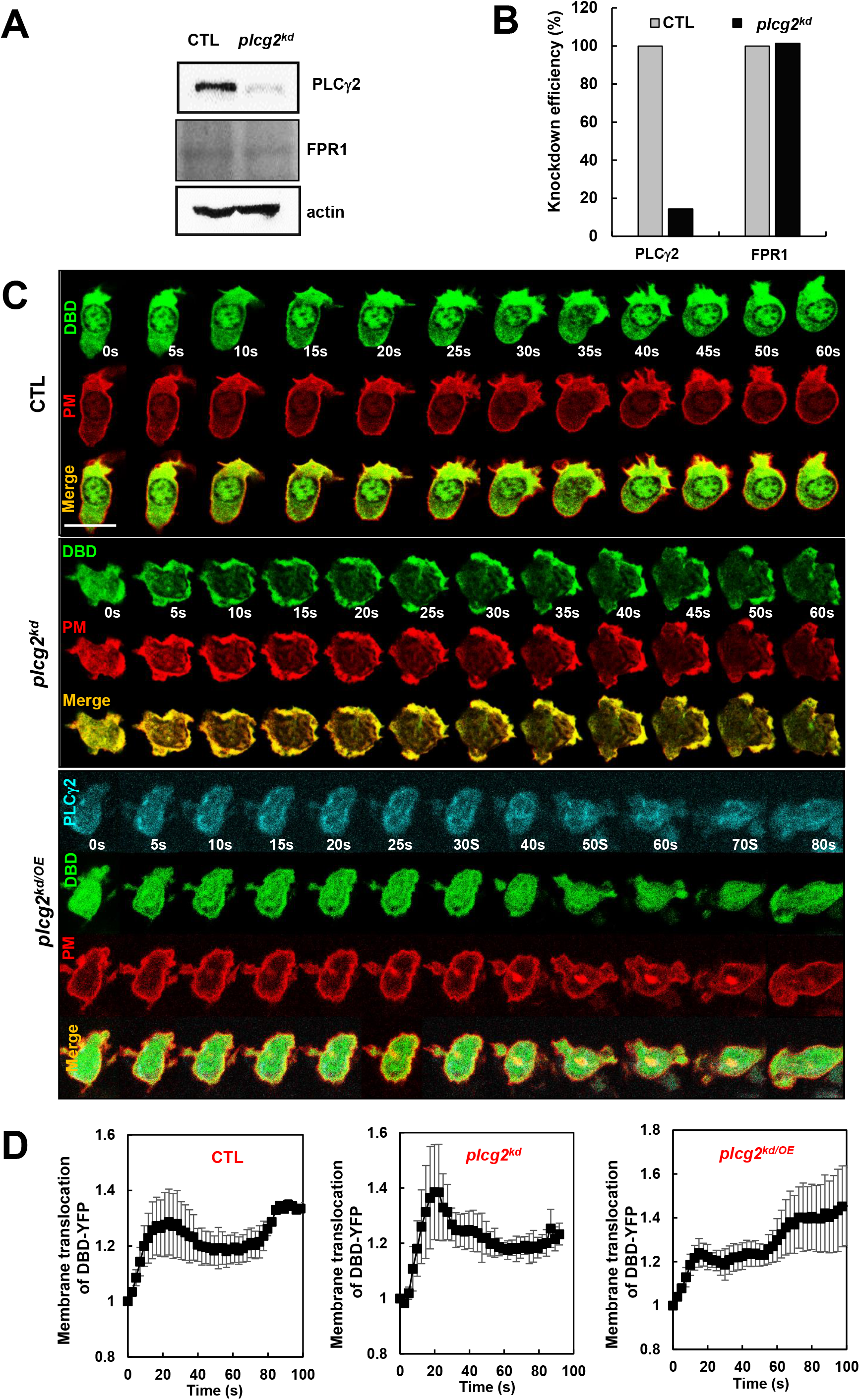
PLCγ2 is an essential part of GPCR-mediated PLC signaling during polarization after the initial response. (A) Expression of *plcg2* in HL60 cells transfected with non-specific (CTL) or *plcg2*-specific (*plcg2^kd^*) shRNA virus particles. PLCγ2 was detected by antibodies recognizing human PLCγ2. Actin was detected as a loading control; fMLP receptor 1 FPR1 was also detected in both CTL and *plcg2^kd^* cells. (B) Quantification of PLCγ2 and FPR1 expression in both CTL and *plcg2^kd^* cells shown in (A). The expression levels of PLCγ2 and FPR1 in CTL cells were normalized to 1. (C) Montage shows fMLP-induced diacylglycerol (DAG) production in CTL, *plcg2^kd^,* and *plcg2*^*kd*/OE^ cells by the membrane translocation of a DAG probe, DAG-binding domain tagged with YFP (DBD-YFP), in response to fMLP stimulation using fluorescence microscopy. Cells expressing DAG-binding domain (DBD) (green) and a PM marker (CAAX-mCherry, red) were stimulated with fMLP at a final concentration of 100 nM right after 0 s. CTL (Video S5) and *plcg2^kd^* (Video S6) cells expressing DAG-binding domain (DBD) (green) and a PM marker (CAAX-mCherry, red) are shown. A *plcg2^kd^*^/OE^ cell (Video S7), which is a *plcg2^kd^* cell expressing PLCγ2 tagged with cerulean (blue), DBD-YFP (green), and a PM marker (CAAX-mCherry, red) is also shown. (D) Quantification of DAG production by the membrane translocation of DAG-receptor in CTL, *plcg2^kd^,* and *plcg2*^*kd*/OE^ cells is shown. Mean ± SD is shown; n= 4, 4 or 5 for CTL, *plcg2^kd^,* and *plcg2*^*kd*/OE^ cells, respectively.

### Impaired calcium responses in *plcg2^kd^* cells

IP3 is a product of PLC activation and initiates calcium release from endoplasmic reticulum (ER) stores followed by a later calcium influx into neutrophils (Clemens and Lowell, 2015). It has been shown that calcium responses in diverse cell types play critical roles in cell migration and chemotaxis (Evans and Falke, 2007; Tsai and Meyer, 2012; Wei et al., 2009). However, chemoattractant-mediated calcium responses in neutrophils, especially in chemotaxing cells, are not well-characterized. Hence, we determined the fMLP-induced calcium response in both CTL and *plcg2^kd^* cells by monitoring the intensity change of a calcium indicator, Fluo-4, using fluorescence microscopy (Fig. 3A). fMLP stimulation induced robust, prolonged and/or multiple calcium responses in CTL cells (Video S8, left), while the same fMLP stimulation induced a similarly robust, but transient, single calcium response in *plcg2^kd^* cells (Video S8, right). The calcium responses of individual CTL and *plcg2^kd^* cells are shown in Fig. S4A-S4B, respectively. By quantitively measuring the intensity change of Fluo-4, we confirmed that both CTL and *plcg2^kd^* cells showed a similar amplitude of calcium response (Fig. 3B), but the duration of calcium response is significantly longer in CTL cells (Fig. 3C). We found that more than 40 % of CTL cells displayed either prolonged or multiple calcium responses, whereas less than 10 % of *plcg2^kd^* cells did so (Fig. S4C). To account for the variation in calcium indicator staining, we normalized calcium response by the equation of (It-I0)/(Imax-I0), I0 and Imax are the Fluo-4 intensity at time 0 s and at maximum response, respectively. Thus, the normalized intensity of Fluo-4 in the cells at time 0 s (I0) is 0 and the intensity at maximum response (Imax) is 1. This normalization confirms that the intracellular calcium-concentration increase in CTL cells lasted significantly longer than that in *plcg2^kd^* cells (Fig. 3D). To understand the role of PLCγ2 in calcium responses in chemotaxing cells, we further monitored calcium response in chemotaxing CTL and *plcg2^kd^* cells experiencing a 1 μM fMLP gradient (Fig. 3E). We found that chemotaxing CTL cells displayed prolonged or often continuous calcium response during continuous chemotaxis toward the source of a chemoattractant gradient (Video S9, left), while *plcg2^kd^* cells showed significantly weaker, sporadic calcium responses at a significantly lower level, and impaired chemotaxis (Video S9, right). Larger views with more cells and more time points are shown in Figure S5. These results together indicate that PLCγ2 activation plays an essential role in maintaining a continuous calcium response during chemotaxis.

**Figure 3.**
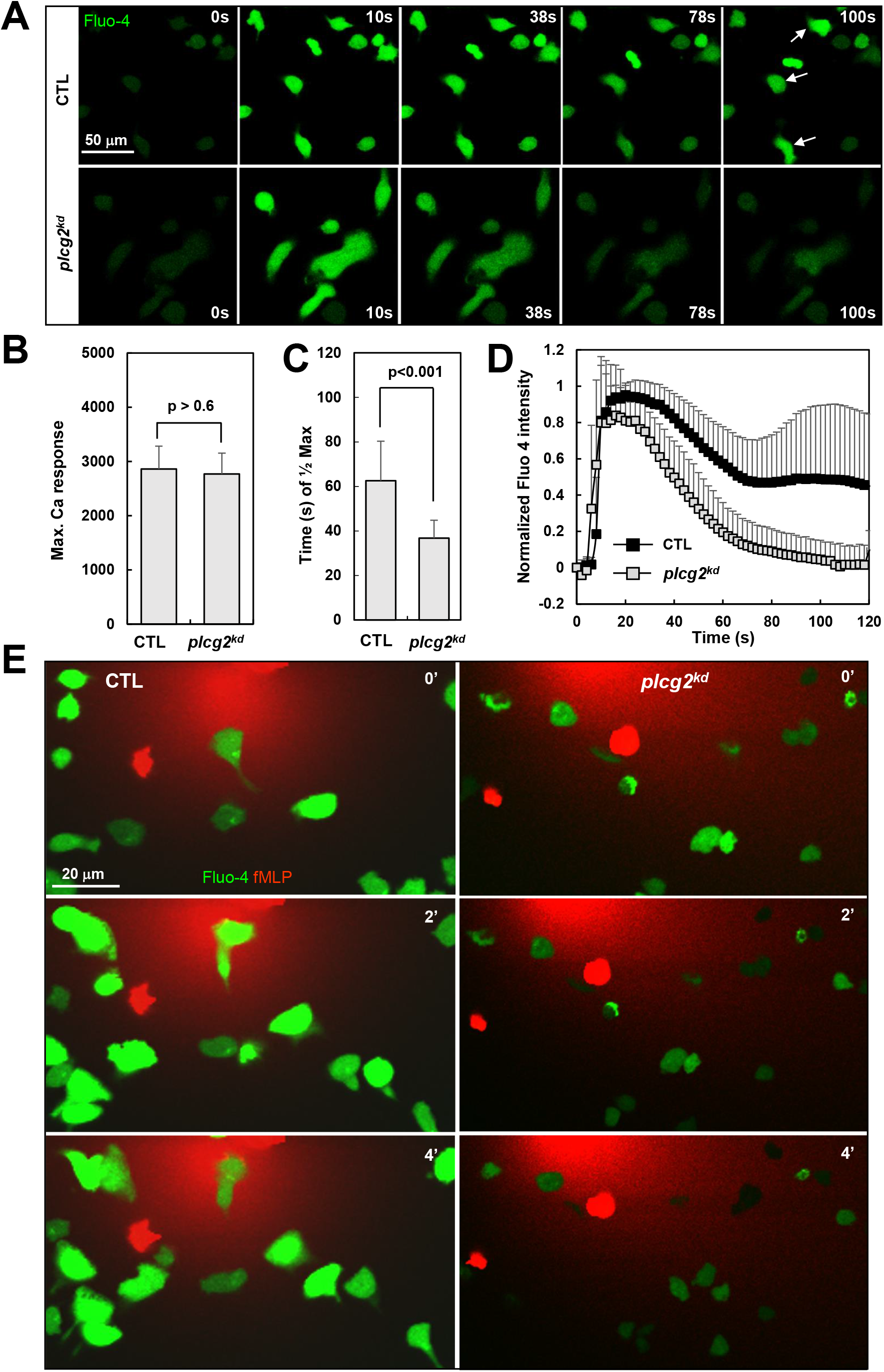
Impaired dynamics of calcium response in *plcg2^kd^* cells. (A) Montage shows 1 μM fMLP-induced calcium response in CTL and *plcg2^kd^* cells. Cells were stained with the fluorescent calcium indicator Fluo-4 (green) and stimulated with 1 μM fMLP at time 0 s. See Video S8, in which the left is CTL cells and the right is *plcg2^kd^* cells. Arrows indicate cells displaying prolonged or multiple calcium responses. (B) Mean ± SD of the maximum calcium response in CTL and *plcg2^kd^* cells. (C) The duration of calcium response was measured as the time required to reach half intensity of maximum response (T1/2). For B and C, mean ± SD is shown, and n = 23 or 22 for CTL or *plcg2^kd^* cells, respectively. The results of Student’s *t*-test are shown. (D) Quantitative measurement of calcium response as fluorescence increase of Fluo-4 in response to chemoattractant fMLP stimulation. To normalize the response, we normalized the lowest intensity of Fluo-4 to 0 and the maximum intensity of calcium response to 1 by the formula, (It-I0)/(Imax-I0). (E) Calcium response in chemotaxing CTL and *plcg2^kd^* cells. Cells were stained with Fluo-4 (green) 30 min prior to the experiment and exposed to a 100 nM fMLP gradient (red). To visualize the gradient, fMLP was mixed with the fluorescent dye Alexa 633 (red). More cells with a larger view at more time points are shown in Fig. S5 and Video S9.

### fMLP stimulation triggers an increased Ras activation in *plcg2^kd^* cells

Proper calcium signaling plays critical roles in the activation and distribution of key components critical for cell migration. Recently, we found that mammalian neutrophils highly express CAPRI, a <u>C</u>alcium-<u>p</u>romoted <u>R</u>as <u>i</u>nactivator, to mediate chemoattractant-induced Ras deactivation for neutrophil chemotaxis, a process that requires a proper chemoattractant-induced calcium response (xuehua xu, 2020). To investigate the effect of the decreased calcium response on the membrane targeting of CAPRI, we monitored the dynamic cellular localization of GFP-tagged CAPRI (green) and a PM marker (red) in both CTL and *plcg2^kd^* cells in response to fMLP stimulation using fluorescence microscopy (Fig. 4A). We found that CAPRI-GFP translocated from the cytoplasm to the PM and colocalized with the PM marker in CTL cells stimulated with 1 μM fMLP (Fig. 4A, upper panel, and Vide S10). However, the same fMLP stimulation triggered significantly less membrane translocation of CAPRI in *plcg2^kd^* cells (Fig. 4A, lower panel, and Video S11). The quantitative measurement of CAPRI membrane translocation from multiple CTL and *plcg2^kd^* cells confirmed the above observation (Fig. 4B). We then monitored Ras activation in the cells by the membrane translocation of an active Ras probe RBD-RFP (active Ras binding domain of human Raf1 tagged with RFP, red) and a PM marker (C1A-C1A-YFP, green) in the cells (Fig. 4C and Fig. S3A) (Xu et al., 2016). We found that in response to 1 μM fMLP stimulation, RBD-RFP translocated from the cytoplasm to the PM and colocalized with the PM marker at the entire cell periphery, then returned to the cytoplasm followed by a second translocation to the protruding sites of CTL cells (Fig. 4C, upper panel, and Video S12). However, the same fMLP stimulation induced robust, persistent membrane translocation of RBD-RFP to the continuously expanding ruffles of *plcg2^kd^* cells (Fig. 4C, lower panel, and Video S13). The quantitative measurement of RBD-RFP membrane translocation showed a typical biphasic Ras activation in CTL cells and a significantly increased Ras activation in *plcg2^kd^* cells (Fig. 4D). These results together indicate that the altered calcium dynamics result in less membrane translocation of CAPRI and consequent elevated Ras activation in *plcg2^kd^* cells.

**Figure 4.**
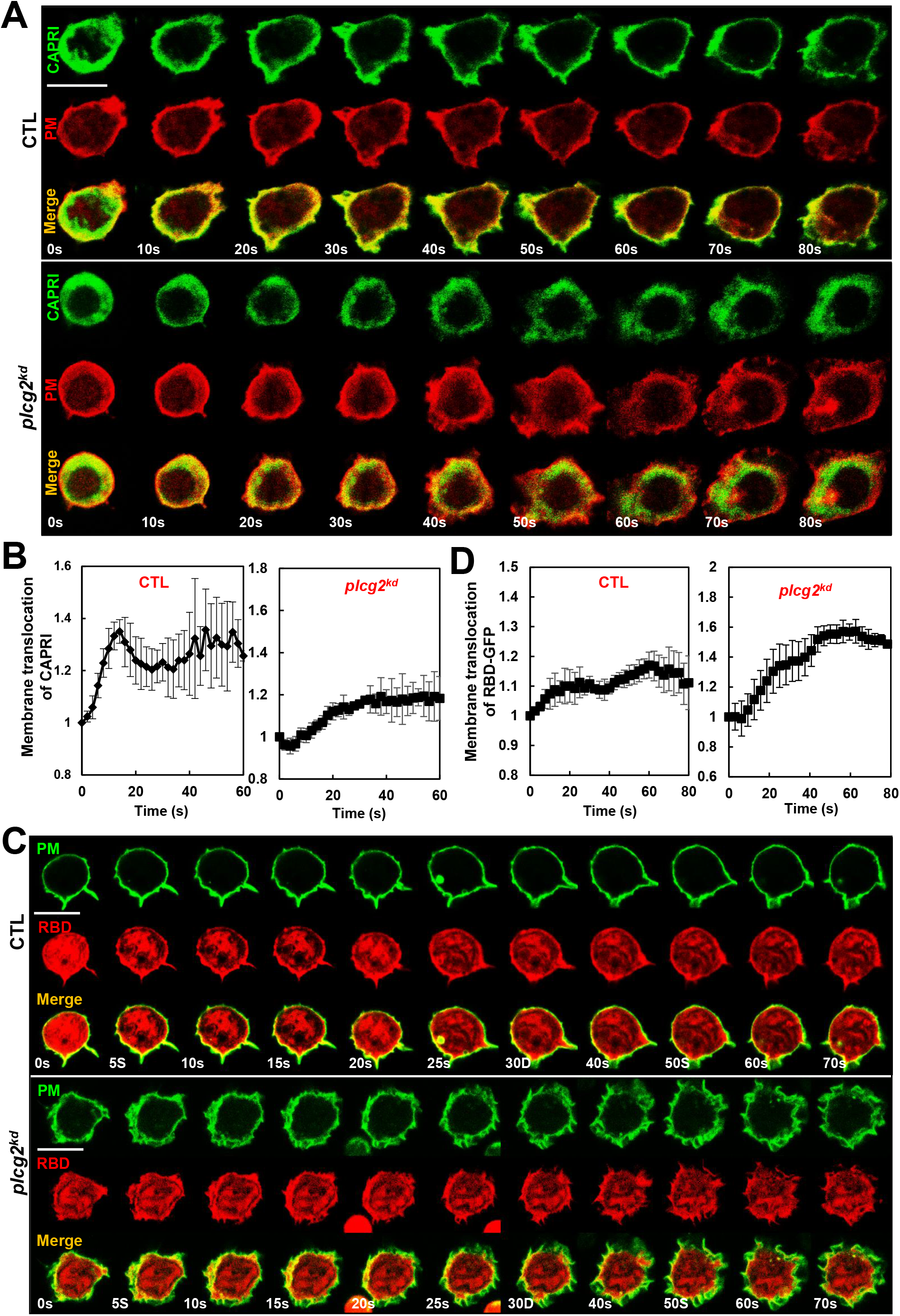
fMLP stimulation induces an increased Ras activation in *plcg2^kd^* cells. (A) Montage shows fMLP-induced membrane translocation of CAPRI in CTL and *plcg2^kd^* cells. CTL cells expressing CAPRI-GFP (green) and a PM marker (CAAX-mCherry, red) were stimulated with fMLP at time 0 s. Scale bar = 10 μm. See Video S10 (CTL) and Video S11 (*plcg2^kd^*) for a complete response of the cells. (B) Quantitative measurement of CAPRI-GFP membrane translocation in CTL and *plcg2^kd^* cells in response to fMLP stimulation. Mean ± SD is shown; n= 4 or 5 for CTL or *plcg2^kd^* cells, respectively. (C) Montage shows fMLP-induced membrane translocation of active Ras probe RBD-RFP in CTL and *plcg2^kd^* cells. Cells expressing RBD-RFP (red) and a PM marker (C1A-C1A-YFP, green) were stimulated with fMLP at time 0 s. Scale bar = 10 μm. See Video S12 (CTL) and S13 (*plcg2^kd^*) for a complete set of cell responses. (D) Quantitative measurement of Ras activation by the RBD-RFP membrane translocation in CTL and *plcg2^kd^* cells in response to fMLP stimulation. Mean ± SD is shown; n= 4 or 4 for CTL or *plcg2^kd^* cells, respectively.

### fMLP stimulation triggers an elevated PI3K activation in *plcg2^kd^* cells

PI_3_Kγ, a direct effector of Ras, synthesizes lipid phosphatidylinositol (3,4,5)-trisphosphate (PtdIns(3,4,5)*P*3, PIP_3_) and recruits and activates the PIP_3_-binding serine/threonine kinase Akt in neutrophils (Li et al., 2000; Suire et al., 2006; Suire et al., 2012). To examine the consequence of the increased Ras activation in *plcg2^kd^* cells, we monitored fMLP-induced PIP_3_ production using a biosensor, PH-GFP (PH-domain of human AKT tagged with GFP, green), by fluorescence microscopy (Fig. 5A) (Servant et al., 2000). We found that 1 μM fMLP triggered a robust translocation of PH-GFP to the entire PM followed by a partial return to the cytosol and a gradual accumulation in the protrusion site of CTL cells (Fig. 5A, upper panel, and Video S14). The same stimulation induced a stronger and persistent translocation of PH-GFP all around the continuously ruffling periphery of *plcg2^kd^* cells (Fig. 5A, lower panel, and Video S15), indicating a hyperactivation of PI3K in cells lacking PLCγ2. Quantitative measurement of membrane translocation of PH-GFP in both multiple CTL and *plcg2^kd^* cells confirmed the above observation (Fig. 5B).

**Figure 5.**
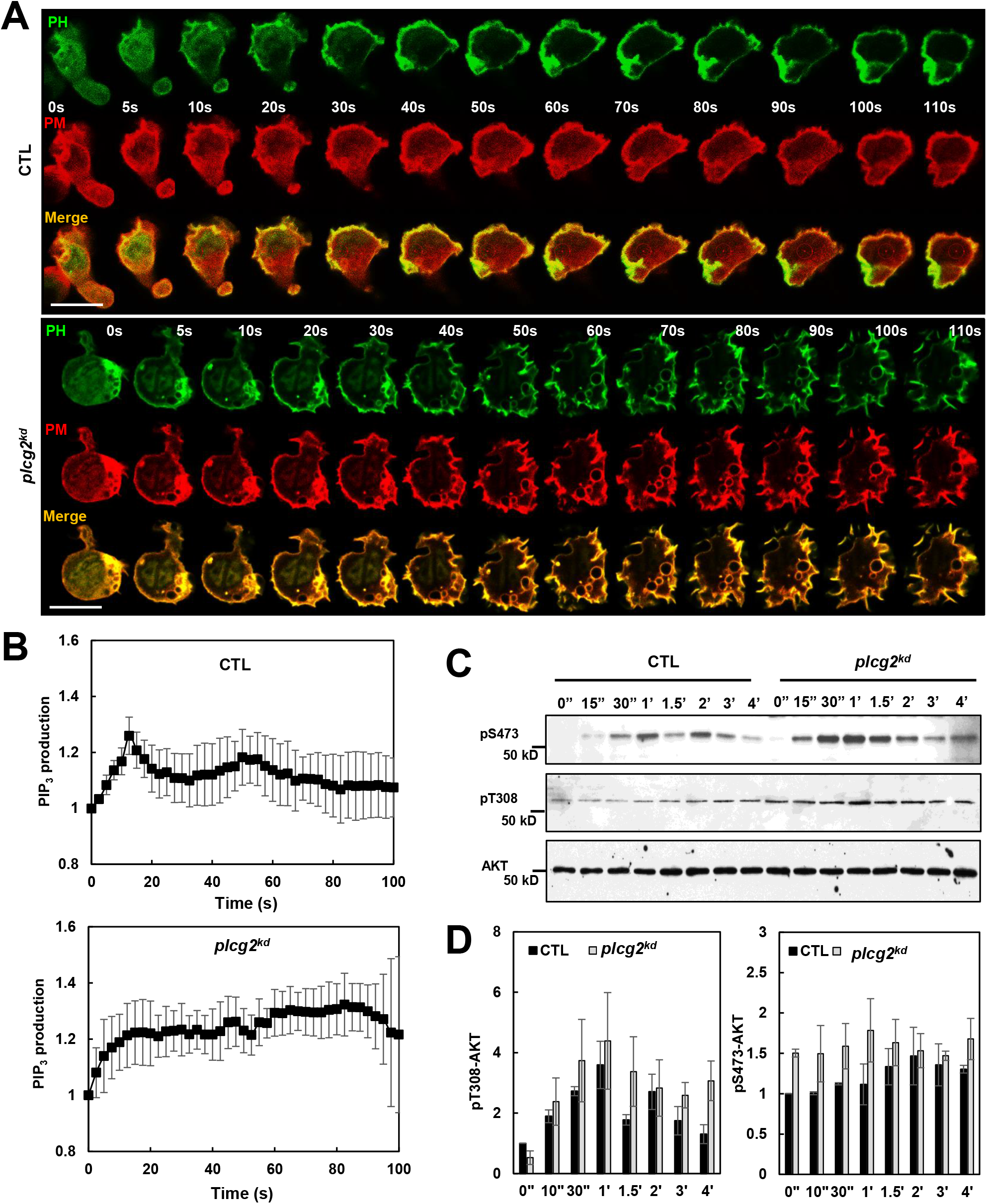
fMLP stimulation triggers an elevated PI3K activation in *plcg2^kd^* cells. (A) Montage shows fMLP-induced PIP_3_ production in CTL and *plcg2^kd^* cells by the membrane translocation of PIP_3_ biosensor PH-GFP. Cells expressing PH-GFP (green) and a PM marker (CAAX-mCherry, red) were stimulated with fMLP at time 0 s. Scale bar = 10 μm. See Video S14 (CTL) and S15 (*plcg2^kd^*) for complete sets of cell responses. (B) Quantitative measurement of PIP_3_ production by the membrane translocation of PH-GFP in CTL and *plcg2^kd^* cells in response to fMLP stimulation. Mean ± SD is shown; n = 3 or 5 for CTL or *plcg2^kd^* cells, respectively. (C) Phosphorylation of AKT on T308 and S473 in CTL and *plcg2^kd^* cells in response to fMLP stimulation detected by western blotting. Cells were stimulated with 1 μM fMLP at time 0 s. Aliquots of the cells at indicated time points were lysed and subjected to western blot detection of the phosphorylated and the total AKT using the indicated antibodies. (D) Quantification of the phosphorylation of AKT at T308 and S473 in (C). Mean ± SD from three independent experiments is shown. The intensity ratio of the phosphorylated and the total AKT in CTL cells at time 0 s was normalized to 1.

fMLP-induced Akt activation is through phosphorylation on threonine 308 (T308) and serine 473 (S473) in a PI3K-dependent manner (Burelout et al., 2007). Thus, we determined fMLP-induced phosphorylation of Akt on these two residues in both CTL and *plcg2^kd^* cells (Fig. 5C). As expected, 1 μM fMLP stimulation induced phosphorylation of Akt on T308 and S473 in CTL cells: a gradually increasing phosphorylation on T308 and a transient, biphasic phosphorylation on S473, consistent with previous reports (Burelout et al., 2007; Cai et al., 2010). In resting *plcg2^kd^* cells, we had detected higher phosphorylation on T308. Stimulation with 1 μM fMLP induced a quicker, stronger phosphorylation on both T308 and S473 followed by a gradual dephosphorylation on both residues (Fig. 5D). Consistent with the above result, an upregulated Akt phosphorylation on both residues induced by fMLP was also observed in *plcb2/3* double knockout (*plcb2/3^-/-^*) murine neutrophils (Tang et al., 2011), suggesting a calcium-dependent regulation of Ras activation in murine neutrophils as well. In conclusion, these results indicate that impaired calcium signaling results in elevated activation of Ras/PI3K and their effectors in human neutrophils.

### Elevated phosphorylation of GSK3 increases cofilin activation in *plcg2^kd^* cells in response to fMLP stimulation

Human neutrophils require a dynamic, well-tuned remodeling of cytoskeleton through actin polymerization and depolymerization in the leading front during chemotaxis. Cofilin is an actin depolymerization factor (ADF) and is essential for depolymerization of F-actin during cell migration (Mizuno, 2013). Cofilin activity is regulated mainly through a phosphorylation event: phosphorylation on Ser3 inhibits its actin binding, severing, and depolymerizing activities; and dephosphorylation on Ser3 by slingshot proteins (SSHs) reactivates it. GSK3 is the first known kinase in murine neutrophils that is phosphorylated by PKC and Akt and deactivates the cofilin phosphatase slingshot 2 (SSH2) to facilitate a dynamic actin polymerization in neutrophils (Tang et al., 2011). Hence, we determined fMLP-induced phosphorylation of GSK3α/3β in both CTL and *plcg2^kd^* cells (Fig. 6A). GSK3 is non-phosphorylated in resting cells, and 1 μM fMLP stimulation induced phosphorylation of GSK3 in CTL cells, consistent with a previous report (Tang et al., 2011). The same fMLP stimulation triggered significantly stronger and more prolonged phosphorylation of GSK3 in *plcg2^kd^* cells (Fig. 6B). Increased phosphorylation of GSK3 is consistent with the elevated activation of Ras/PI3K in these cells (Fig. 4 and Fig. 5). Next, to determine the consequence of the altered phosphorylation profiles of GSK3, we examined cofilin dephosphorylation/activation in both CTL and *plcg2^kd^* cells upon fMLP stimulation (Fig. 6C). As previously reported (Tang et al., 2011; Xu et al., 2015b), fMLP stimulation transiently decreased the amount of phosphorylated cofilin in CTL cells. The amount of phosphorylated cofilin returned to the pre-stimulation state at around 90 s in CTL cells. The same 1 μM fMLP stimulation, however, decreased phosphorylation of cofilin in *plcg2^hi^* cells (Fig. 6D). These results indicate that elevated phosphorylation of GSK3 increases dephosphorylation and activation of cofilin in *plcg2^kd^* cells.

**Figure 6.**
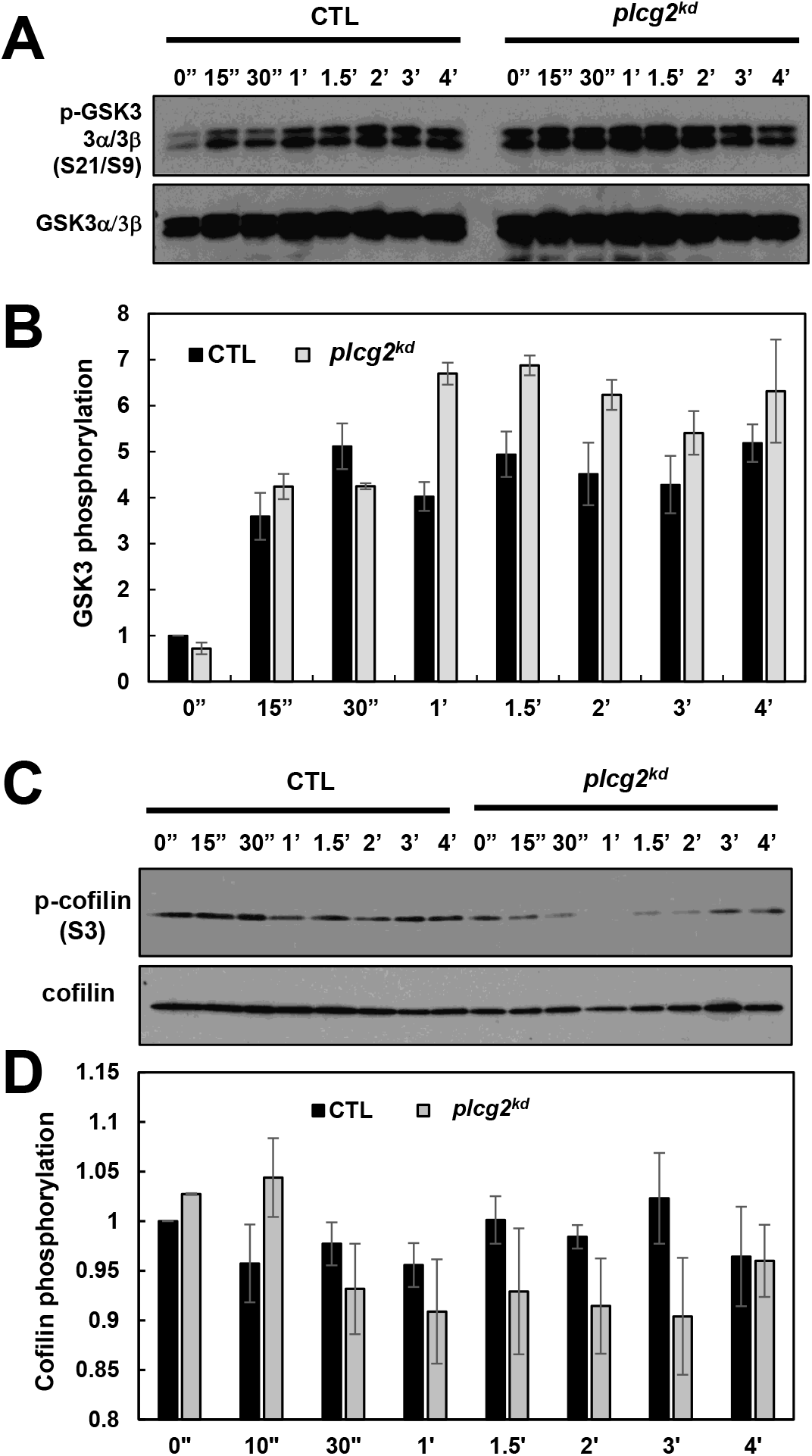
Elevated phosphorylation of GSK3 increases the phosphorylation of cofilin in *plcg2^kd^* cells in response to fMLP stimulation. (A) Phosphorylation of GSK3 in CTL and *plcg2^kd^* cells in response to fMLP stimulation detected by western blotting. Cells were stimulated with 1 μM fMLP at time 0 s. Aliquots of the cells at the indicated time points were lysed and subjected to western blot detection of the phosphorylated and the total GSK3α/3β using their specific antibodies. (B) Quantification of the phosphorylation of GSK3 in (**A)**. Mean ± SD from three independent experiments is shown. The intensity ratio of the phosphorylated and total proteins in CTL cells at time 0 s was normalized to 1. (C) Cofilin activation/dephosphorylation in CTL and *plcg2^kd^* cells in response to fMLP stimulation. Cells were stimulated with 1 μM fMLP at time 0 s. Aliquots of the cells at indicated time points were lysed and subjected to western blot detection of phosphorylated and total cofilin using their specific antibodies. (D) Quantification of cofilin dephosphorylation in CTL and *plcg2^kd^* cells in response to 1 μM fMLP stimulation. Mean ± SD from three independent experiments is shown. The ratio of phosphorylated cofilin versus total cofilin in CTL cells at time 0 s was normalized to 1.

### Impaired dynamics of fMLP-induced actin polymerization, polarization, and migration during chemotaxis in *plcg2^kd^* cells

The chemoattractant GPCR/G-protein signaling regulates multiple signaling pathways to control the actin cytoskeleton dynamics that drive cell migration. To evaluate the role of PLCγ2 in chemoattractant GPCR-mediated actin assembly in neutrophils, we determined fMLP-mediated polymerization of actin in CTL and *plcg2^kd^* cells using a centrifugation assay of actin filaments (F-actin) (Fig. 7A). In CTL cells, 10 μM fMLP stimulation induced a transient polymerization of actin. In *plcg2^kd^* cells, the same stimulation triggered prolonged polymerization (Fig. 7A-7B). To understand the temporospatial dynamics of actin polymerization, we next monitored actin polymerization using the membrane translocation of an F-actin filament probe, F-tractin-GFP, in live cells by fluorescence microscopy (Yi et al., 2012). We found that, upon fMLP stimulation at 2 s, F-tractin-GFP translocated to the cell cortex at around 10 to 20 s, mostly returned to the cytosol at about 30 s, and then translocated to the leading front again at around 40 s in CTL cells (Fig. 7C, upper panel, and Video S16), indicating a biphasic actin polymerization (Cai et al., 2010; Charest et al., 2010). In *plcg2^k^* cells, the same fMLP stimulation triggered persistent membrane translocation of F-tractin to the cell membrane, indicating a continuous actin polymerization in the cells (Fig. 7C, lower panel, and Video S17). We further monitored the distribution of F-actin in migrating cells toward a 1 μM fMLP gradient using an F-actin probe (SiR-actin) (Fig. 7D). Resting CTL cells displayed polarized morphology. Upon exposure to a 1 μM fMLP gradient, CTL cells quickly reoriented and migrated toward the source of 1 μM fMLP while displaying even better polarized morphology: a protruding leading front (pseudopod) and a contracting trailing edge (uropod) (Video S17, left). Resting *plcg2^kd^* cells displayed less polarized, rounder morphology. In response to an fMLP gradient at 1 μM fMLP, *plcg2^kd^* cells often migrated continuously in their original direction with significantly impaired polarization and decreased speed over time (Video S17, right). We further quantified the polarization of CTL and *plcg2^kd^* cells by the ratio of width versus length of the cells and confirmed that *plcg2^kd^* cells displayed a significant defect in cellular polarization in a fMLP gradient compare to CTL cells (Fig. 7E).

**Figure 7.**
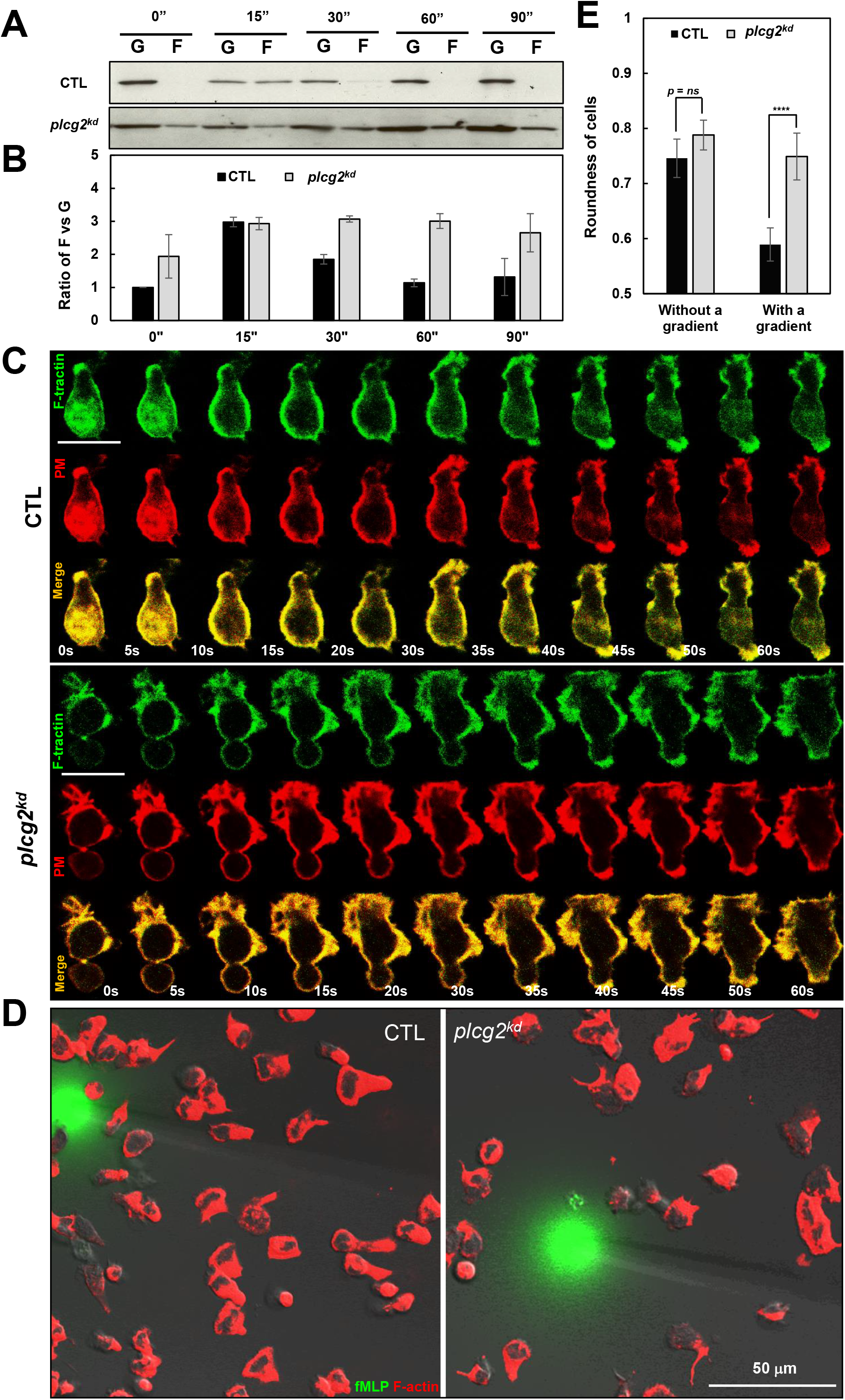
Impaired dynamics of actin polymerization in *plcg2^kd^* cells. (A) The amount of globular (G) and filamentous (F) actin in CTL and *plcg2^kd^* cells was determined by a centrifugation assay of F-actin. Cells were stimulated with 10 μM fMLP at time 0 s and aliquots of cells at the indicated time points were analyzed. (B) Normalized quantitative densitometry of the F/G-actin ratio in the CTL and of *plcg2^kd^* cells in (A). Mean ± SD from three independent experiments is shown. The F/G ratio of CTL cells at 0 s was normalized to 1. (C) Montage shows the membrane translocation of F-actin probe (GFP-tagged F-tractin) in CTL and *plcg2^kd^* cells upon fMLP stimulation. Cells expressing F-tractin GFP (green) and a PM marker (CAAX-mCherry, red) were stimulated with fMLP at time 0 s. Scale bar = 10 μm. See Video S16 (CTL) and S17 (*plcg2^kd^*) for a complete set of cell responses. (D) F-actin distribution in chemotaxing CTL and *plcg2^kd^* cells in a 1 μM fMLP gradient. Cells were stained with the actin filament probe SiR-actin (red) and chemotaxed in a fMLP gradient (green) for about 3 min. To visualize the gradient, fMLP was mixed with Alexa 488 (green, 1 μg/ml). Scale bar = 50 μm. (E) The graph shows the roundness of CTL and *plcg2^kd^* cells shown in (C). Roundness (%) as the parameter of the polarity of a cell was measured by the ratio of the width to the length of a cell. That is, the roundness for a circle is 1 and for a line is 0. Mean ± SD is shown. N = 33 for both CTL and *plcg2^kd^* cells. The *p* values of Student’s */*-test are indicated as *ns* (not significant,*p* > 0.1), * (*p* < 0.1), ** (*p* < 0.01), or *** (*p* < 0.001), or **** (*p* < 0.0001).

We then examined the chemotaxis behavior of CTL and *plcg2^kd^* cells in gradients generated from the source of either 1 μM fMLP or 100 ng/mL IL8 using an *EZ-TaxiScan* chemotaxis assay (Fig. 8A). We found that, as compared to CTL cells, *plcg2^kd^* cells displayed clear chemotaxis defects measured as four parameters of chemotaxis behavior: total path length, directionality, speed, and polarity in both fMLP and IL8 gradients (Fig. 8B). All of the above results indicate that GPCR-mediated PLCγ2 activation is required for GPCR-mediated chemotaxis of human neutrophils.

**Figure 8.**
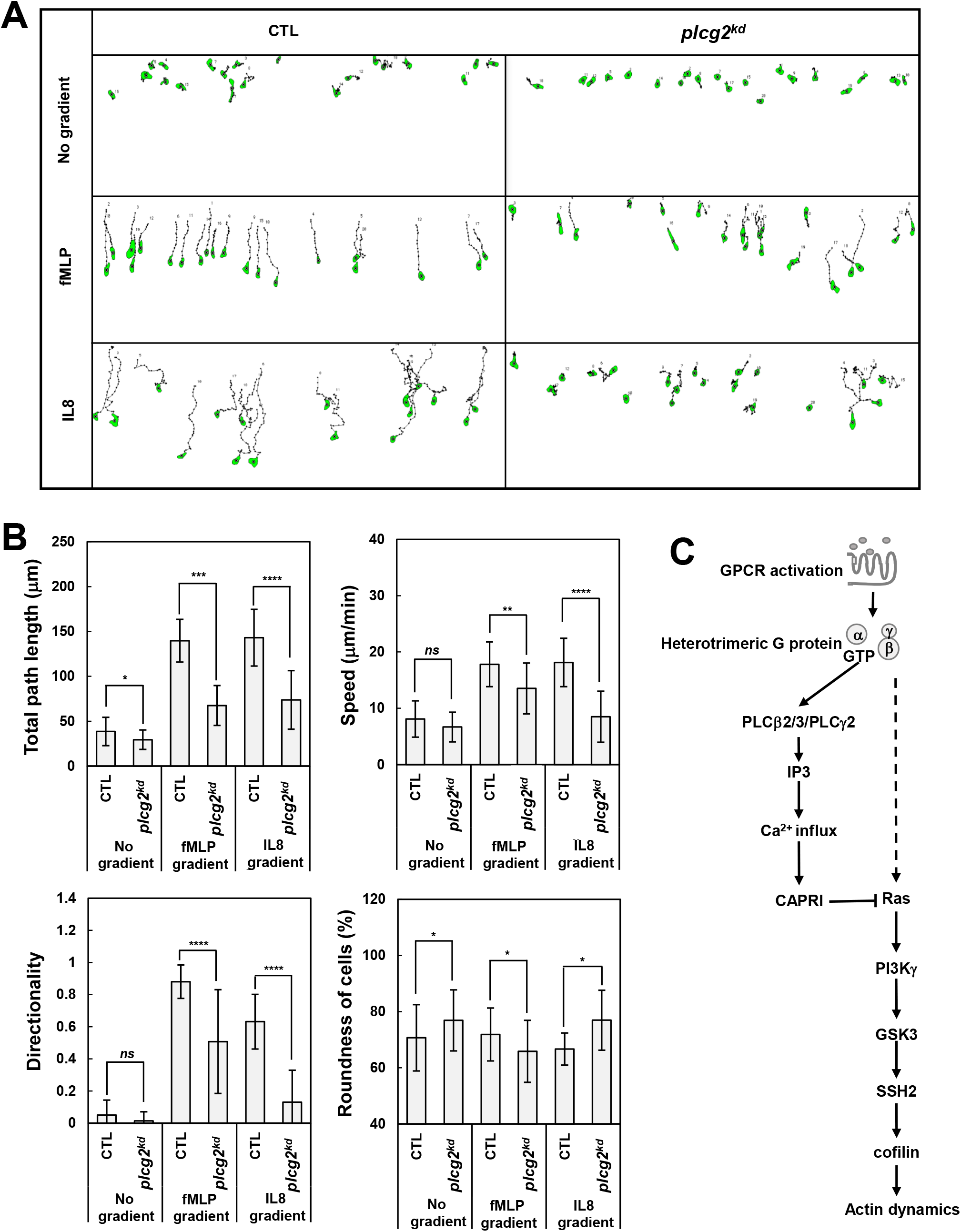
*plcg2^kd^* cells display defects in chemoattractant-mediated chemotaxis. (A) Montage shows the traveling path of CTL and *plcg2^kd^* cells experiencing no gradient (NG) or gradients generated from a 1 μM fMLP or 1 μg/mL IL8 source. See Video S18 for cell migration. (B) Chemotaxis behaviors measured from (A) and described as four parameters: directionality, where 0 represents random movement and 1 represents straight movement to the micropipette; speed, defined as the distance that a cell’s centroid moves as a function of time; total path length, the total distance a cell has traveled; and roundness, an indication of cell polarization, where 0 represents a straight line, perfect polarization; and 100% represents a circle, non-polarization. Twenty cells in each group were analyzed. The time lengths of data in fMLP and IL8 gradients are 3.5 min and 7.25 min, respectively. Mean ± SD (standard deviation) is shown. The *p* values of Student’s Ltest are indicated as *ns* (not significant,*p* > 0.1), * (*p* < 0.1), ** (*p* < 0.01), or *** (*p* < 0.001), or **** (*p* < 0.0001). (C) Scheme shows the GPCR-mediated PLCγ2 activation and signaling pathways to regulate actin dynamics during chemotaxis of neutrophils.

### Gain-of-function mutant of PLCγ2 improves chemotaxis capability of neutrophils at low temperature

Cold treatment or injury induces an acute immune response that involves increased production of multiple cytokines and chemokines at the lesion site (Koedel et al., 2007). One main characteristic of cold-induced urticaria is increased neutrophil infiltration, a process closely related to neutrophil chemotaxis. PLCγ2-associated antibody deficiency and immune dysregulation (PLAID) patients, who bear gain-of-function mutations of PLCγ2 caused by genomic deletions in PLCγ2 and suffer from cold-induced urticaria (Ombrello et al., 2012). Aderibigbe et al further reported enhanced activation of neutrophils and monocytes in PLAID and APLAID patients, consistent with the essential role of PLCγ2 in the activation of neutrophils or monocytes (Aderibigbe et al., 2015). Surprisingly, the neutrophils from PLAID patients display decreased chemotaxis, though chemotaxis is an essential process for neutrophil infiltration. Therefore, we first examined the chemotaxis behavior of HL60 cells expressing either empty vector, or vector encoding either WT or deletion mutant of Δ686, which is a gain-of-function mutant and equivalent to the deletion mutation of PLAID patients (Wang et al., 2014). We transiently transfected chemotactic HL60 cells with plasmids encoding chimeric proteins of green fluorescent protein (GFP) alone or GFP-fused to the C terminus of either wildtype (WT) PLCγ2 or mutant Δ686, and then we purified GFP-expressing cells by fluorescence-activated cell sorting (FACS). Using these GFP expressing cells, we confirmed that cells expressing Δ686 exhibited decreased chemotaxis compared to those expressing either empty vector or WT-PLCγ2 at 37 °C, consistent with the previous report (Fig. 9 and Video S19) (Aderibigbe et al., 2015). We next examined the chemotaxing capability of the above cells at two lower temperatures. At 30 °C, cells expressing either Δ686 or WT of PLCγ2 chemotaxed similarly but better than the cells expressing empty vector. At 28 °C, cells expressing Δ686 chemotaxed significantly better than those expressing WT, which is better than the cells expressing empty vector. The above result indicates that the gain-of-function mutant of PLCγ2 improves the chemotaxis capability of cells at low temperature, while it worsens chemotaxis at body temperature.

**Figure 9.**
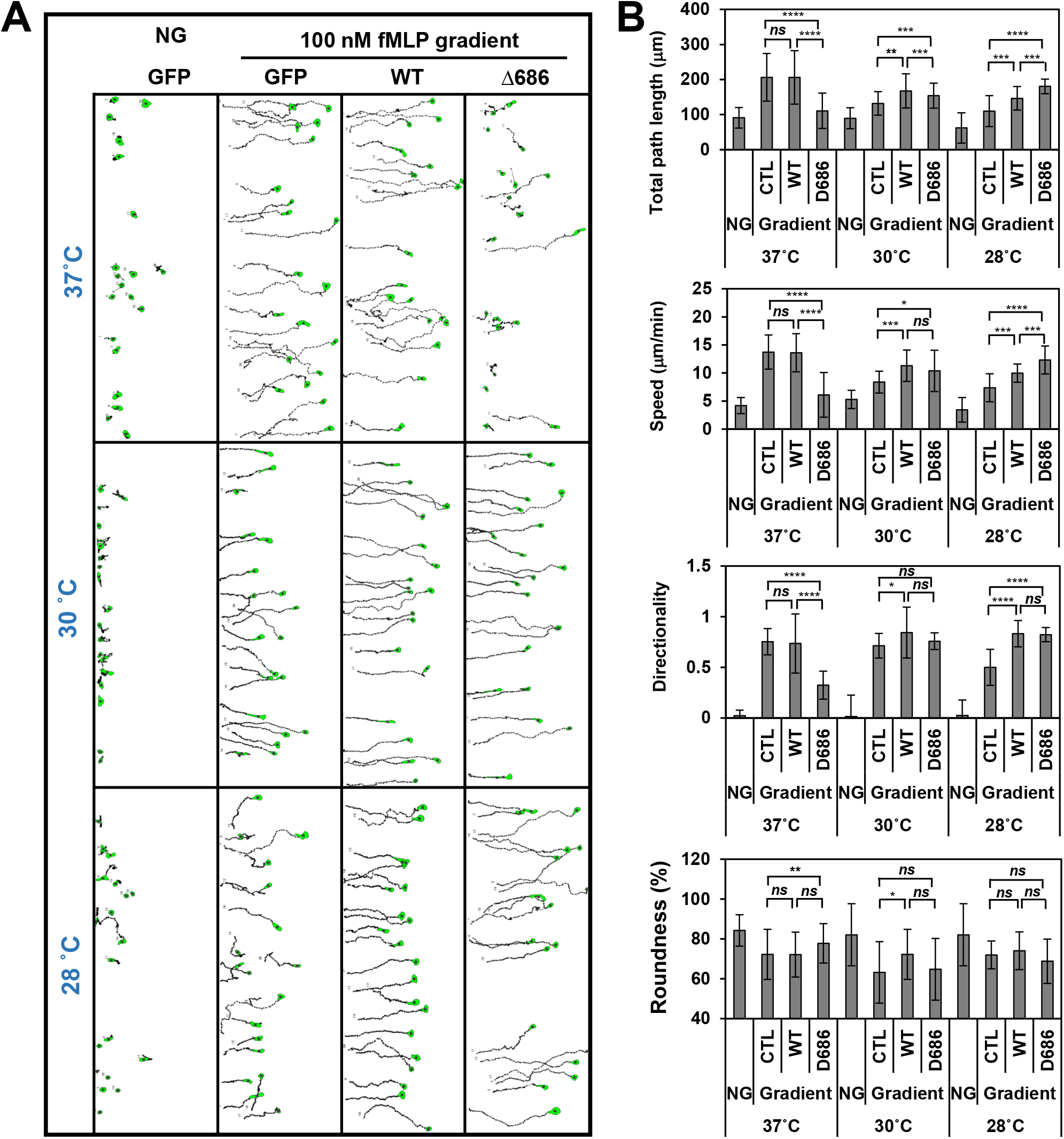
Cells expressing gain-of-function mutant of PLCγ2 (D686) display better chemotaxis than cells expressing either empty or wild-type PLCγ2. (A) Montage shows the traveling path of CTL cells experiencing no gradient (NG) or cells expressing either GFP vector, or GFP-tagged with wild-type (WT) or truncated mutant D686 of PLCγ2 and experiencing fMLP gradients generated from a 100 nM fMLP source. See Video S19 for cell migration. See Video S19 for the cell migration under the indicated conditions. (B) Chemotaxis behaviors measured from (A) and described as four parameters: directionality, where 0 represents random movement and 1 represents straight movement to the micropipette; speed, defined as the distance that a cell’s centroid moves as a function of time; total path length, the total distance a cell has traveled; and roundness, an indication of cell polarization, where 0 represents a straight line, perfect polarization; and 100% represents a circle, non-polarization. Twenty cells in each group were analyzed. The time lengths of data shown are 14 min. Mean ± SD (standard deviation) is shown. The *p* values of Student’s *t*-test are indicated as *ns* (not significant *p* > 0.1), * (*p* < 0.1), ** (*p* < 0.01), *** (*p* < 0.001), or **** (*p* < 0.0001).

## DISCUSSION

It is believed that chemoattractants/GPCRs activate β phospholipase C (PLCβ) and that receptor tyrosine kinases (RTKs) activate γ phospholipase C (PLCγ). In the present study, we show that chemoattractant/GPCR-mediated membrane recruitment and subsequent activation of PLCγ2 constitutes an essential part of GPCR-mediated PLC signaling for polarization and migration during neutrophil chemotaxis (Fig. 8C). Importantly, cells expressing the gain-of-function mutant of PLCγ2 exhibit better chemotaxis than cells expressing wild-type (WT) at low temperature, providing a molecular mechanism for the initiation of dysregulated recruitment and activation of neutrophils during cold-induced urticaria of PLAID patients.

In mammals, the 13 PLC isoforms are divided into 6 subgroups: -β, -γ, -δ, -ε, -, and -η (Suh et al., 2008). Mammalian neutrophils dominantly express three isoforms: PLC-β2, -β3, and -γ2. Free Gα or Gβγ of heterotrimeric G protein directly activates PLCβ, either in response to chemoattractant stimulation or in an *in vitro* assay (Camps et al., 1992; Jiang et al., 1997; Jiang et al., 1994; Li et al., 2000). PLCβ2, -β3 and -γ2 share a common domain structure (Fig. S6A). They all contain a PH, an EF, a C2 domain, and an X-Y catalytic domain. We found that chemoattractant stimulation induces robust membrane translocation of only PLCγ2, and not PLCβ2/β3 (Fig. S6B). In a detailed comparison of the PH domains of PLCβ2/b3/γ2 and PIP_3_- or PIP2-binding PH domains, we found that the PH domain of PLCγ2 possesses the highest homology with PIP_3_-binding PH domain, and the PH domains of PLCβ2 and -β3 have better homology with those PIP2-binding PH domains (Fig. S6C). In addition, the C2 domain of PLCγ2 contains all key residues essential for calcium binding, but the domains of PLCβ2 and -β3 do not (Fig. S6D). The above explains the fact that the membrane-translocation capability of PLCγ2 is decreased in the mutants that lack either the PH or the C2 domain (Fig. 1), consistent with the previous reports that PLCγ2 membrane translocation requires calcium response and PIP_3_ production (Falasca et al., 1998; Nishida et al., 2003; Xu et al., 2015b). Importantly, in response to uniform chemoattract stimulation, *plcg2^kd^* cells exhibited similar levels of the initial PLC activation but lacked the secondary DAG production and calcium response (Fig. 2 and Fig. 3). The altered dynamics of DAG production and calcium response in *plcg2^kd^* cells, in response to either uniform or gradient stimulations, indicate that PLCγ2-mediated PLC signaling play essential roles in repolarization and continuous migration in a chemoattractant gradient. These results together suggest that chemoattractant/GPCR-mediated PIP_3_ production and PLCβ-mediated [Ca^2+^] increase act together to recruit and activate PLCγ2 to achieve PLC signaling for polarization and continuous migration during chemotaxis after the initial response.

Chemoattractant-mediated activation of protein kinase C (PKC) is mediated by PLC signaling and plays critical roles in the chemotaxis and other functions of neutrophils (Li et al., 2000; Liu et al., 2014; Tang et al., 2011; Xu et al., 2015b). Neutrophils express PKCα, -βl, -βII, and -δll (Bertram and Ley, 2011). PKC activation is mainly regulated through phosphorylation/dephosphorylation at three conserved phosphorylation sites known as the activation-loop (A-loop), the turn motif (TM), and the hydrophobic motif (HM) (Fig. S7A-S7B) (Freeley et al., 2011). Murine neutrophils with a double deficiency of PLCβ2/3 (*plcb2^-/-^ plcb3^-/-^*) display significantly decreased fMLP-induced initial DAG and IP3 production, and activation of PKC isoforms (Li et al., 2000). In contrast to the above, we detected altered dynamics of DAG production and calcium response in HL60 cells lacking PLCγ2 (*plcg2^kd^*) (Fig. 2 and Fig. 3). We also found that fMLP stimulation induced an unexpected higher level of PKC phosphorylation in *plcg2^kd^* cells (Fig. S8). Consistent with the above, we detected highly phosphorylated PKCα/β on (T638/T642) and PKCα(T497), the two residues that critically control stability and accessibility for the dephosphorylation and inactivation of PKCs (Bertram and Ley, 2011; Bornancin and Parker, 1996). The highly phosphorylated status of PKC on these two residues might hinder the dephosphorylation process of PKCs, providing an explanation for the elevated phosphorylation of PKCs in *plcg2^kd^* cells. However, the connection between PLCγ2 deficiency and the phosphorylation state of PKCs needs further investigation.

### Limitation of the study

The result shown in **Figure 9** is significant. We acknowledge that the necessity and impact of this assay using neutrophils derived from PLAID patients are far beyond the work done in human cell lines. Lack of access to PLAID patients and different phenotypes observed in mouse models prevented us from repeating and confirming our conclusion. As a result, the data are presented as a proof of concept that future investigations and applications will be built upon.

## Supporting information

Supplementary informations

## ACKNOWLEDGMENTS

We thank Silver Poland and Wenxiang Sung for providing the bone marrow of mice for the work and Dustin Parsons for reading the manuscript and providing helpful suggestions. This work was supported by the NIH Intramural Fund from the National Institute of Allergy and Infectious Diseases, National Institutes of Health.

## MATERIALS AND METHODS

### Cell culture and differentiation

HL60-CXCR2 cells were obtained from Ann Richmond of Vanderbilt University. HL60 cells were purchased from ATCC Inc. and cultured, maintained, and differentiated as previously reported (Xu et al., 2015a). Briefly, cells were maintained in RPMI 1640 culture medium [RPMI 1640 medium with 20% (v/v) fetal bovine serum and 25 mM HEPES (Quality Biological, Inc. Gaithersburg, MD)]. HL60 cells were differentiated in RPMI 1640 culture medium containing 1.3% DMSO for 5 days before the experiments. The cells were incubated at 37°C in a humidified 5% CO2 atmosphere.

### Establishment of stable PLCγ2 knockdown cell line

On Day 1, 2.5 × 10^5^ cells were seeded in a 6-well plate. On Day 2, the culture medium in the 6-well plate was replaced by 1 ml of infection cocktail (RPMI 1640 culture medium containing 5 μg/ml polybrene). Virus particles encoding a *plcg2-specific* or a non-specific shRNA (Santa Cruz Biotechnology, Santa Cruz, CA) were added to the cells. Six hours later, the infection cocktail of each well was replaced by RPMI 1640 culture medium without polybrene. On Day 3, puromycin (Sigma Aldrich, St. Louis, MO) was added to the cells at a final concentration of 0.3 μg/ml. The culture medium was changed every 2-3 days until the infected cells proliferated enough for the experiments.

### Reagents and antibodies

f-Met-Leu-Phe (fMLP) and DMSO were from Sigma Aldrich (St. Louis, MO); α-PLCγ2, -p(phospho)-PLCγ2(Y759), -AKT, -p-AKT(T308), -p-AKT(T473), -Erk, -p-Erk, -cofilin, -p-cofilin(S3), -GSK3α/3β, -p-GSK3α/3β (S21/S9), -actin, and -FPR1 antibodies were from Cell Signaling Technology (Beverly, MA); HRP-conjugated anti-mouse or anti-rabbit IgG was from Jackson ImmunoResearch (West Grove, PA); and all tissue culture reagents were from Invitrogen (Carlsbad, CA).

### Plasmids and transfection of cells

The DNA vectors of GFP-tagged PLCγ2 and its mutants, and turboGFP (tGFP)-tagged human CAPRI and control tGFP alone were from OriGene Inc. (Gaithersburg, MD) and will be deposited to Addgene.com. Active Ras sensor (active Ras binding domain of human Raf1 tagged with mRFP, RBD-RFP), diacylglycerol probes (diacylglycerol-binding domain of PKCβll tagged with YFP, DBD-YFP), and PM markers (C1AC1A-YFP, CAAX-mCherry, and Memcerulean) were from Addgene (Cambridge, MA). F-actin sensor F-tractin-GFP was obtained from John Hammer (Yi et al., 2012). The transfection procedure was as previously described (Xu et al., 2016). Briefly, 2 × 10^6^ cells were centrifuged at 100 × *g* for 10 min and resuspended in a mixture of 80 μL nucleofection solution V and 20 μL supplement I at room temperature. Six micrograms of plasmid DNA encoding the cDNA of the desired proteins was used for a single transfection reaction using program T-019 on the Amaxa Nucleofector II (Lonza, MD).

### Immunoblotting of fMLP-mediated phosphorylation of proteins

The differentiated cells were starved for 1 hour prior to the experiments. Cells were stimulated with fMLP at the final concentration of 1 μM. Aliquots of cells at the indicated time points were directly added to the same volume of 2X SDS loading buffer (SLB) and subjected to SDS-PAGE and western blot detection.

### Calcium response

Cells were incubated with 100 ng/ml Fluo-4 (Invitrogen, Carlsbad, CA) at 37°C for 30 min, washed with RPMI 1640 medium with 25 mM HEPES twice to remove the unstained Fluo-4, and then subjected to the experiments.

#### Imaging and data processing

Cells were plated and allowed to adhere to the cover glass of a 4-well or a 1-well chamber (Nalge Nunc International, Naperville, IL) precoated with Fibronectin (Sigma Aldrich, Saint Louis, MO) for 10 min, and then covered with RPMI 1640 medium with 10% FBS and 25 mM HEPES. For confocal microscopy, cells were imaged using a Carl Zeiss Laser Scanning Microscope Zen 780 (Carl Zeiss, Thornwood, NY) with a Plan-Apochromat 60x/1.4 Oil DIC M27 objective. For the uniform-stimulation experiment of membrane translocation assays, the stimuli were directly delivered to the cells as previously described (Xu et al., 2016). The micropipette chemotaxis assay was done as previously reported (Xu et al., 2009). To establish a gradient, a Femtotip micropipette is filled with 30 μl of solution containing chemoattractant and a desirable fluorescent dye (either Alexa 488 or Alexa 633) and then attached to the FemtoJet microinjector and micromanipulator. The output pressure of the FemtoJet is set at Pc = 70 hPa to release a constant and tiny volume of the mixture into a one-well chamber that is filled with 6 mL of buffer and cells. Under these conditions, a gradient can be established within 100 μm from the tip of the micropipette and is usually stable for more than 1 h. Images were processed and analyzed with Zen Black software. Images were further processed in Adobe Photoshop (Adobe Systems, San Jose, CA). The membrane translocation of the indicated protein was measured by the depletion of the interested protein in the cytoplasm. The data obtained were further analyzed with Microsoft Office Excel (Redmond, WA). For quantitative analysis of membrane translocation dynamics of the indicated molecules, the cytosolic depletion of the indicated molecule was measured. Regions of interest (ROIs) in the cytoplasm (avoiding the nucleus area as much as possible) were within the cells throughout the time period of the measurements. The periphery of the cells was marked by the membrane markers. For data analysis, to normalize the effect of photobleaching during data acquisition, the intensity of ROIs in the cytoplasm was first divided by the intensity of whole cells at each given time point. To normalize the effect of morphological change during the time period, the above resulting data were divided by the intensity of ROIs in the PM marker channel. Lastly, the resulting data were divided by that at time 0 s; consequently, the relative intensity of any cells at time 0 s became 1. The graph of mean ± SD is shown.

#### Actin polymerization assay

The protocol of G and F actin measurement mostly followed the instructions of the “G-actin /F-actin *In Vivo* Assay Kit” from Cytoskeleton Inc. (Denver, CO) (Xu et al., 2017). Briefly, 100 μl aliquots of cells at the indicated time points before and after 10 μM fMLP stimulation were mixed with 500 μl LAS2 buffer containing 1X LAS1 buffer (provided in the kit), 10 mM ATP, and 1X proteinase inhibitor cocktail (Roche Life Science, Indianapolis, IN). The mixture of samples was homogenized using a 200 μl pipet tip and incubated at 37°C for 10 min. Next, the samples were centrifuged at 350 × *g* for 10 min to pellet unbroken cells. A total of 400 μl supernatant of each sample was transferred to a centrifugation tube and centrifuged at 100,000 × *g* at 4°C for 1 hour. This step is to pellet the F-actin and leave the G-actin in the supernatant. Supernatants were transferred to prelabeled 1.8 ml centrifugation tubes and mixed with 400 μl SDS loading buffer. To the pellet, 400 μl actin depolymerization buffer was added and incubated on ice for 1 hour to allow actin depolymerization to occur. Then, 400 μl SDS loading buffer was added to each pellet sample. 10 μl of each sample was subjected to SDS-PAGE and western blot to detect and analyze G- and F-actin amounts at given time points for the cell lines.

#### SiR-actin staining

The differentiated HL60 cells were washed with RPMI1640 medium once and transferred into one well of a 24-well plate. The actin filament probe SiR-actin (Cytoskeleton, Inc.) and verapamil were added to the cells at a final concentration of 1 μM and 10 μM, respectively, and incubated for 1 hour at 37 °C. The cells were then washed with RPMI 1640 starvation medium with 10 μM verapamil, added to the fibronectin pre-coated one-well chambers, and incubated at 37 °C for 3 hours prior to imaging.

#### TAXIScan chemotaxis assay and data analysis

The procedure was as previously reported (Wen et al., 2016). Briefly, differentiated cells were loaded onto fibronectin-coated 4-μm EZ-TAXIScan chambers. The chemoattractants at the indicated concentrations were added to the other side of the well across the terrace that the cells chemotax through. The cells migrated for 30 min at 37 °C. Images were taken at 30-s intervals. For chemotaxis parameter measurements, 20 cells in each group were analyzed with DIAS software (Wessels et al., 1998). The bar graphs of chemotaxis parameters in mean and SD were plotted with Microsoft Office Excel (Redmond, WA).

## ACKNOWLEDGMENTS

We thank Silver Poland and Wenxiang Sung for providing the bone marrow of mice for the work. We thank Heawon Song for providing the plasmid of PLCγ2-D686. This work was supported by the NIH intramural fund from the National Institute of Allergy and Infectious Diseases, National Institutes of Health.

## AUTHOR CONTRIBUTIONS

Conceptualization: X.X.; Investigation: X. X., X.W., S.B., A.M., D.P.; Data analysis: X. X., X.W., S.B.; Writing – Original draft, X.X., Review & Editing, X.X., T.J., X.W., S.B., A.M., D.P.; Funding acquisition, T.J.

## CONFLICTS OF INTEREST

The authors declare that they have no conflict of interest.

## Abbreviations used in this paper

GPCR: G protein-coupled receptor
PLC: phospholipase C
PLCß2/β3: β2/β3 phospholipase C
PLCγ2: γ2 phospholipase C
WT: wild-type
DAG: diacylglycerol
DBD: DAG-binding domain of PKC
PKC: protein kinase C
RBD: Ras-binding domain of human Raf1
CAPRI: calcium-promoted Ras inactivator
SSH: slingshot protein
PKD1: protein kinase D1
PH-domain: pleckstrin-homologue domain.

## Notes

### Competing Interest Statement

The authors have declared no competing interest.

## REFERENCES

Aderibigbe, O.M., D.L. Priel, C.C. Lee, M.J. Ombrello, V.H. Prajapati, M.G. Liang, J.J. Lyons, D.B. Kuhns, E.W. Cowen, and J.D. Milner. 2015. Distinct Cutaneous Manifestations and Cold-Induced Leukocyte Activation Associated With PLCG2 Mutations. JAMA Dermatol. 151:627–634.

Bertram, A., and K. Ley. 2011. Protein kinase C isoforms in neutrophil adhesion and activation. Arch Immunol Ther Exp (Warsz). 59:79–87.

Bornancin, F., and P.J. Parker. 1996. Phosphorylation of threonine 638 critically controls the dephosphorylation and inactivation of protein kinase Calpha. Curr Biol. 6:1114–1123.

Burelout, C., P.H. Naccache, and S.G. Bourgoin. 2007. Dissociation between the translocation and the activation of Akt in fMLP-stimulated human neutrophils--effect of prostaglandin E2. J Leukoc Biol. 81:1523–1534.

Cai, H., S. Das, Y. Kamimura, Y. Long, C.A. Parent, and P.N. Devreotes. 2010. Ras-mediated activation of the TORC2-PKB pathway is critical for chemotaxis. J Cell Biol. 190:233–245.

Camps, M., A. Carozzi, P. Schnabel, A. Scheer, P.J. Parker, and P. Gierschik. 1992. Isozyme-selective stimulation of phospholipase C-beta 2 by G protein beta gamma-subunits. Nature. 360:684–686.

Charest, P.G., Z. Shen, A. Lakoduk, A.T. Sasaki, S.P. Briggs, and R.A. Firtel. 2010. A Ras signaling complex controls the RasC-TORC2 pathway and directed cell migration. Dev Cell. 18:737–749.

Clemens, R.A., and C.A. Lowell. 2015. Store-operated calcium signaling in neutrophils. J Leukoc Biol. 98:497–502.

Evans, J.H., and J.J. Falke. 2007. Ca2+ influx is an essential component of the positive-feedback loop that maintains leading-edge structure and activity in macrophages. Proc Natl Acad Sci U S A. 104:16176–16181.

Falasca, M., S.K. Logan, V.P. Lehto, G. Baccante, M.A. Lemmon, and J. Schlessinger. 1998. Activation of phospholipase C gamma by PI 3-kinase-induced PH domain-mediated membrane targeting. The EMBO journal. 17:414–422.

Freeley, M., D. Kelleher, and A. Long. 2011. Regulation of Protein Kinase C function by phosphorylation on conserved and non-conserved sites. Cell Signal. 23:753–762.

Gallegos, L.L., M.T. Kunkel, and A.C. Newton. 2006. Targeting protein kinase C activity reporter to discrete intracellular regions reveals spatiotemporal differences in agonist-dependent signaling. The Journal of biological chemistry. 281:30947–30956.

Jakus, Z., E. Simon, D. Frommhold, M. Sperandio, and A. Mocsai. 2009. Critical role of phospholipase Cgamma2 in integrin and Fc receptor-mediated neutrophil functions and the effector phase of autoimmune arthritis. The Journal of experimental medicine. 206:577–593.

Jiang, H., Y. Kuang, Y. Wu, W. Xie, M.I. Simon, and D. Wu. 1997. Roles of phospholipase C beta2 in chemoattractant-elicited responses. Proc Natl Acad Sci U S A. 94:7971–7975.

Jiang, H., D. Wu, and M.I. Simon. 1994. Activation of phospholipase C beta 4 by heterotrimeric GTP-binding proteins. The Journal of biological chemistry. 269:7593–7596.

Koedel, U., U.M. Merbt, C. Schmidt, B. Angele, B. Popp, H. Wagner, H.W. Pfister, and C.J. Kirschning. 2007. Acute brain injury triggers MyD88-dependent, TLR2/4-independent inflammatory responses. The American journal of pathology. 171:200–213.

Kolaczkowska, E., and P. Kubes. 2013. Neutrophil recruitment and function in health and inflammation. Nat Rev Immunol. 13:159–175.

Koss, H., T.D. Bunney, S. Behjati, and M. Katan. 2014. Dysfunction of phospholipase Cgamma in immune disorders and cancer. Trends Biochem Sci. 39:603–611.

Li, Z., H. Jiang, W. Xie, Z. Zhang, A.V. Smrcka, and D. Wu. 2000. Roles of PLC-beta2 and-beta3 and PI3Kgamma in chemoattractant-mediated signal transduction. Science. 287:1046–1049.

Liew, P.X., and P. Kubes. 2019. The Neutrophil’s Role During Health and Disease. Physiol Rev. 99:1223–1248.

Liu, L., D. Gritz, and C.A. Parent. 2014. PKCbetaII ACTS DOWNSTREAM OF CHEMOATTRACTANT RECEPTORS AND mTORC2 TO REGULATE cAMP PRODUCTION AND MYOSIN II ACTIVITY IN NEUTROPHILS. Molecular biology of the cell.

Mizuno, K. 2013. Signaling mechanisms and functional roles of cofilin phosphorylation and dephosphorylation. Cell Signal. 25:457–469.

Nathan, C. 2006. Neutrophils and immunity: challenges and opportunities. Nature reviews. Immunology. 6:173–182.

Nishida, M., K. Sugimoto, Y. Hara, E. Mori, T. Morii, T. Kurosaki, and Y. Mori. 2003. Amplification of receptor signalling by Ca2+ entry-mediated translocation and activation of PLCgamma2 in B lymphocytes. The EMBO journal. 22:4677–4688.

Nourshargh, S., and R. Alon. 2014. Leukocyte migration into inflamed tissues. Immunity. 41:694–707.

Ombrello, M.J., E.F. Remmers, G. Sun, A.F. Freeman, S. Datta, P. Torabi-Parizi, N. Subramanian, T.D. Bunney, R.W. Baxendale, M.S. Martins, N. Romberg, H. Komarow, I. Aksentijevich, H.S. Kim, J. Ho, G. Cruse, M.Y. Jung, A.M. Gilfillan, D.D. Metcalfe, C. Nelson, M. O’Brien, L. Wisch, K. Stone, D.C. Douek, C. Gandhi, A.A. Wanderer, H. Lee, S.F. Nelson, K.V. Shianna, E.T. Cirulli, D.B. Goldstein, E.O. Long, S. Moir, E. Meffre, S.M. Holland, D.L. Kastner, M. Katan, H.M. Hoffman, and J.D. Milner. 2012. Cold urticaria, immunodeficiency, and autoimmunity related to PLCG2 deletions. The New England journal of medicine. 366:330–338.

Park, D., D.Y. Jhon, C.W. Lee, K.H. Lee, and S.G. Rhee. 1993. Activation of phospholipase C isozymes by G protein beta gamma subunits. The Journal of biological chemistry. 268:4573–4576.

Ramirez, R.N., N.C. El-Ali, M.A. Mager, D. Wyman, A. Conesa, and A. Mortazavi. 2017. Dynamic Gene Regulatory Networks of Human Myeloid Differentiation. Cell Syst. 4:416–429 e413.

Rincon, E., B.L. Rocha-Gregg, and S.R. Collins. 2018. A map of gene expression in neutrophil-like cell lines. BMC Genomics. 19:573.

Servant, G., O.D. Weiner, P. Herzmark, T. Balla, J.W. Sedat, and H.R. Bourne. 2000. Polarization of chemoattractant receptor signaling during neutrophil chemotaxis. Science. 287:1037–1040.

Suh, P.G., J.I. Park, L. Manzoli, L. Cocco, J.C. Peak, M. Katan, K. Fukami, T. Kataoka, S. Yun, and S.H. Ryu. 2008. Multiple roles of phosphoinositide-specific phospholipase C isozymes. BMB Rep. 41:415–434.

Suire, S., A.M. Condliffe, G.J. Ferguson, C.D. Ellson, H. Guillou, K. Davidson, H. Welch, J. Coadwell, M. Turner, E.R. Chilvers, P.T. Hawkins, and L. Stephens. 2006. Gbetagammas and the Ras binding domain of p110gamma are both important regulators of PI(3)Kgamma signalling in neutrophils. Nature cell biology. 8:1303–1309.

Suire, S., C. Lecureuil, K.E. Anderson, G. Damoulakis, I. Niewczas, K. Davidson, H. Guillou, D. Pan, C. Jonathan, T.H. Phillip, and L. Stephens. 2012. GPCR activation of Ras and PI3Kc in neutrophils depends on PLCb2/b3 and the RasGEF RasGRP4. The EMBO journal. 31:3118–3129.

Tang, W., Y. Zhang, W. Xu, T.K. Harden, J. Sondek, L. Sun, L. Li, and D. Wu. 2011. A PLCbeta/PI3Kgamma-GSK3 signaling pathway regulates cofilin phosphatase slingshot2 and neutrophil polarization and chemotaxis. Dev Cell. 21:1038–1050.

Tsai, F.C., and T. Meyer. 2012. Ca2+ pulses control local cycles of lamellipodia retraction and adhesion along the front of migrating cells. Curr Biol. 22:837–842.

Violin, J.D., J. Zhang, R.Y. Tsien, and A.C. Newton. 2003. A genetically encoded fluorescent reporter reveals oscillatory phosphorylation by protein kinase C. The Journal of cell biology. 161:899–909.

Wang, J., H. Sohn, G. Sun, J.D. Milner, and S.K. Pierce. 2014. The autoinhibitory C-terminal SH2 domain of phospholipase C-gamma2 stabilizes B cell receptor signalosome assembly. Science signaling. 7:ra89.

Wei, C., X. Wang, M. Chen, K. Ouyang, L.S. Song, and H. Cheng. 2009. Calcium flickers steer cell migration. Nature. 457:901–905.

Wen, X., T. Jin, and X. Xu. 2016. Imaging G Protein-coupled Receptor-mediated Chemotaxis and its Signaling Events in Neutrophil-like HL60 Cells. J Vis Exp.

Wessels, D., E. Voss, N. Von Bergen, R. Burns, J. Stites, and D.R. Soll. 1998. A computer-assisted system for reconstructing and interpreting the dynamic three-dimensional relationships of the outer surface, nucleus and pseudopods of crawling cells. Cell Motil Cytoskeleton. 41:225–246.

Xu, X., J.A. Brzostowski, and T. Jin. 2009. Monitoring dynamic GPCR signaling events using fluorescence microscopy, FRET imaging, and single-molecule imaging. Methods Mol Biol. 571:371–383.

Xu, X., N. Gera, H. Li, M. Yun, L. Zhang, Y. Wang, J. Wang, and T. Jin. 2015a. GPCR-Mediated PLCbetagamma/PKCbeta/PKD Signaling Pathway Regulates the Cofilin Phosphatase Slingshot 2 in Neutrophil Chemotaxis. Molecular biology of the cell.

Xu, X., N. Gera, H. Li, M. Yun, L. Zhang, Y. Wang, Q.J. Wang, and T. Jin. 2015b. GPCR-mediated PLCbetagamma/PKCbeta/PKD signaling pathway regulates the cofilin phosphatase slingshot 2 in neutrophil chemotaxis. Molecular biology of the cell. 26:874–886.

Xu, X., X. Wen, D.M. Veltman, I. Keizer-Gunnink, H. Pots, A. Kortholt, and T. Jin. 2017. GPCR-controlled membrane recruitment of negative regulator C2GAP1 locally inhibits Ras signaling for adaptation and long-range chemotaxis. Proc Natl Acad Sci U S A. 114:E10092–E10101.

Xu, X., M. Yun, X. Wen, J. Brzostowski, W. Quan, Q.J. Wang, and T. Jin. 2016. Quantitative Monitoring Spatiotemporal Activation of Ras and PKD1 Using Confocal Fluorescent Microscopy. Methods in molecular biology. 1407:307–323.

xuehua xu, x.w., amer moosa, smit bhimani, tian jin. 2020. Ras Inhibitor CAPRI Enables Neutrophils to Chemotax Through a Higher-Concentration Range of Gradients.

Yi, J., X.S. Wu, T. Crites, and J.A. Hammer, 3rd. 2012. Actin retrograde flow and actomyosin II arc contraction drive receptor cluster dynamics at the immunological synapse in Jurkat T cells. Molecular biology of the cell. 23:834–852.

Zhou, Q., G.S. Lee, J. Brady, S. Datta, M. Katan, A. Sheikh, M.S. Martins, T.D. Bunney, B.H. Santich, S. Moir, D.B. Kuhns, D.A. Long Priel, A. Ombrello, D. Stone, M.J. Ombrello, J. Khan, J.D. Milner, D.L. Kastner, and I. Aksentijevich. 2012. A hypermorphic missense mutation in PLCG2, encoding phospholipase Cgamma2, causes a dominantly inherited autoinflammatory disease with immunodeficiency. Am J Hum Genet. 91:713–720.

