## Supplementary informations for "GPCR-mediated Activation of PLCγ2 Initiates Dysregulated Recruitment of Neutrophils in Cold-induced Urticaria in PLAID Patients"

**SUPPLEMENTARY MATERIALS**

**Supplemental movies:**

**Video S1**. A control (CTL) HL60 cell expressing a plasma membrane marker (CAAX-mCherry, red) and GFP-tagged wild-type (PLCγ2-WT, green) was stimulated with 1 μM fMLP at time 0 s. Scale bar = 10 μm.

**Video S2**. A CTL HL60 cell expressing a plasma membrane marker (CAAX-mCherry, red) and GFP-tagged lipase-dead (PLCγ2-LD, green) was stimulated with 1 μM fMLP at time 0 s. Scale bar = 10 μm.

**Video S3**. A CTL HL60 cell expressing a plasma membrane marker (CAAX-mCherry, red) and GFP-tagged PH domain-deleted mutant of PLCγ2 (PLCγ2-ΔPH, green) was stimulated with 1 μM fMLP at time 0 s. Scale bar = 10 μm.

**Video S4**. A CTL HL60 cell expressing a plasma membrane marker (CAAX-mCherry, red) and GFP-tagged C2 domain-deleted mutant of PLCγ2 (PLCγ2-ΔC2, green) was stimulated with 1 μM fMLP at time 0 s. Scale bar = 10 μm.

**Video S5**. A CTL cell expressing DAG probe, DBD-YFP (green), and plasma membrane (PM) marker, CAAX-RFP (red), was stimulated with fMLP stimulus at 0 s. Scale bar = 10 μm.

**Video S6**. A $plcg2^{kd}$ HL60 cell, in which *plcg2* expression was knocked down and expressing probe, DBD-YFP (green), and plasma membrane (PM) marker, CAAX-RFP (red), was stimulated with fMLP stimulus at 0 s. Scale bar = 10 μm.

**Video S7**. A *plcg2*$^{kd/OE}$ cell expressing DAG probe, DBD-YFP (green), a plasma membrane (PM) marker, CAAX-RFP (red), and/or PLCγ2-cerulean (blue) was stimulated with fMLP stimulus at 0 s. Scale bar = 10 μm.

**Video S8**. Calcium response in CTL (left) or *plcg2*$^{kd}$ (right) cells. Cells were stained with the calcium indicator, Fluo-4, and stimulated with 1 μM fMLP at the beginning of the movies. Scale bar = 50 μm.

**Video S9**. Calcium response in chemotaxing CTL (left) or *plcg2*$^{kd}$ (right) cells in a 100 nM fMLP gradient (red). Cells were stained with the calcium indicator Fluo-4 (green). Scale bar = 50 μm.

**Video S10. A** CTL cell expressing GFP-CAPRI (green) and a PM marker (CAAX-mCherry, red) was stimulated with 1 μM fMLP at 0 s. Scale bar = 10 μm.

**Video S11.** A *plcg2*$^{kd}$ cell expressing GFP-CAPRI (green) and a PM marker (CAAX-mCherry, red) was stimulated with 1 μM fMLP at 0 s. Scale bar = 10 μm.

**Video S12**. A CTL cell expressing active Ras probe (mRFP-RBD, red) and a PM marker (C1A-C1A-YFP, green) was stimulated with 1 μM fMLP at 0 s.. Scale bar = 10 μm.

**Video S13**. A *plcg2*$^{kd}$ cell expressing active Ras probe (mRFP-RBD, red) and a PM marker (C1A-C1A-YFP, green) were stimulated with 1 μM fMLP at 0 s. Scale bar = 10 μm.

**Video S14**. A CTL cell expressing PIP3 probe (PH-GFP, green) and PM marker (CAAX-mCherry, red) was stimulated with 1 μM fMLP at 0 s. Scale bar = 10 μm.

**Video S15**. A *plcg2*$^{kd}$ cell expressing $PIP_3$ probe (PH-GFP, green) and PM marker (CAAX-mCherry, red) was stimulated with 1 μM fMLP at 0 s. Scale bar = 10 μm.

**Video S16**. A CTL cell expressing F-actin probe (Ftractin-GFP, green) and PM marker (CAAX-mCherry, red) was stimulated with 1 μM fMLP at 0 s. Scale bar = 10 μm.

**Video S17**. A *plcg2*$^{kd}$ cell expressing F-actin probe (Ftractin-GFP, green) and PM marker (CAAX-mCherry, red) was stimulated with 1 μM fMLP at 0 s. Scale bar = 10 μm.

**Video S18.** Montaged movie shows the movement of CTL (left) or *plcg2*$^{kd}$ (right) cells experiencing no gradient (top), or gradients generated from 1 μM fMLP (middle) or 1 μg/ml IL8 (bottom) sources.

**Video S19.** Montaged movie shows the movement of CTL cells expressing empty GFP vector (column 1, no gradient), GFP vector (column 2, 100 nM fMLP gradient), wild-type of PLCγ2 (WT) (column 3, 100 nM fMLP gradient), or deletion mutant of 686 (column 4, 100 nM fMLP gradient) from the left to the right at the indicated temperature.

**Supplementary figure legends**

Fig. S1. PLCγ2 is highly expressed in mammalian neutrophils and neutrophil-like (HL60) cells. The expression pattern of PLC isoforms in primary human and mouse neutrophils (**A**) or in human neutrophil-like (HL60) cells (**B**).

Fig. S2. The scheme shows the domain composition of wild-type (WT), lipase-dead (LD), and the truncated (ΔPH and ΔC2) mutants of PLCγ2.

Fig. S3. Three plasma membrane (PM) markers colocalize in HL60 cells before and after chemoattractant fMLP stimulation. (**A**) Colocalization of two membrane markers **(**C1AC1A-YFP and CAAX-mCherry) in HL60 cells. Cell expressing two plasma membrane (PM) markers, C1AC1A-YFP (green) and CAAX-mCherry (red), was stimulated with fMLP at time 0s. During the period of imaging, C1AC1A-YFP colocalizes with in the cells before and after fMLP stimulation. (**B**) Colocalization of two membrane markers **(**mem-cerulean and CAAX-mCherry) in HL60 cells. Cell expressing two plasma membrane (PM) markers, mem-cerulean (cyan) and CAAX-mCherry (red), was stimulated with fMLP at time 0 s. Mem-cerulean colocalizes with CAAX-mCherry before and after fMLP stimulation. Scale bar = 10 μm in **A** and **B**.

Fig. S4. fMLP-induced calcium response in CTL and $plcg2^{kd}$ cells. Calcium responses in the individual CTL (**A**) and $plcg2^{kd}$ (**B**) cells in response to fMLP stimulation. See supplementary Video 8. Calcium response is visualized by the fluorescent calcium indicator Fluo-4. Cells were stained with Fluo-4 (green) 30 min prior to the experiment. CTL and $plcg2^{kd}$ cells were stimulated with 100 nM fMLP at time 8 s or 6 s, respectively. The numbers in **A** and **B** indicates an individual cell in the same single experiment. (**C**) The percentage of CTL or $plcg2^{kd}$ cells that displayed either a single/transient (grey) or prolonged/multiple calcium increase (black). N= 95 or 57 cells in CTL or $plcg2^{kd}$ cells, respectively.

Fig. S5. Calcium response in chemotaxing CTL and $plcg2^{kd}$ cells. Cells were stained with Fluo-4 (green) 30 min prior to the experiment and exposed to a 100 nM fMLP gradient (red). To visualize the gradient, fMLP was mixed with fluorescent dye Alexa 633 (red). See supplemental Video S9, in which the left is CTL cells and the right is $plcg2^{kd}$ cells.

Fig. S6. Chemoattractant stimulation induces a strong membrane translocation of PLCγ2, but not PLCβ2/β3 in HL60 cells. (**A**) The scheme shows the domain composition of PLCβ2, -β3, and -γ2. (**B**) Chemoattractant stimulation induces a strong membrane translocation of PLCγ2, but not PLCβ2/β3 in HL60 cells. HL60 cells were stimulated at time 0 s. Aliquots of the cells at the indicated time points were mechanically lysed using 5 μm filter systems. The membrane fractions of the cells were collected by centrifugation at 10,000 × *g* for 10 min. The pellets were mixed with SDS loading buffer and subject to western blotting detection of the indicated proteins. CSF1R were detected as the loading control of the membrane fraction of the cells. (**C**) Alignments of the PH domains of human PLCβ1/β2/γ2 and two PIP$_3$-binding PH domains of human Akt and BTK in the upper panel; the alignment of the PH domains of PLCβ1/β2/γ2 and two PIP$_2$-binding PH domains of Human PLCδ1 and PLCL1in the lower panel. (**D**) Alignment of the C2 domains of human PLCβ1/β2/γ2 and two calcium-binding C2 domains of human SYT1 and PKCβ1. Sequence data in (C) and (D) were obtained from https://www.ncbi.nlm.nih.gov/pubmed/ and aligned using the ClustalW2 provided from https://www.ebi.ac.uk.

Fig. S7. Conserved residues of protein sequences and phosphorylation sites among protein kinase C (PKC) isoforms. (**A**) The scheme shows the domain composition and phosphorylation sites of PKCα/β, and -γ. (**B**) Conserved phosphorylation sites of PKC isoforms in neutrophils. Sequence data were obtained from https://www.ncbi.nlm.nih.gov/pubmed/ and aligned using the ClustalW2 provided from https://www.ebi.ac.uk.

Fig. S8. Altered phosphorylation dynamics of PKC in *plcg2*$^{kd}$ cells. (**A**) fMLP-induced phosphorylation dynamics of PKC isoforms in CTL and *plcg2*$^{kd}$ cells. Aliquots of cells at the indicated time points before and after 1 μM fMLP stimulation were lysed and subjected to western blot analysis using the indicated antibodies. (**B**) Quantification of PKC isoform phosphorylation upon uniform fMLP stimulation in (**A**). The intensity ratio of the phospho-PKC isoform and total PKC in CTL at time 0 s was normalized to 1. Mean ± SD from three independent experiments is shown.

**Supplementary materials and methods**

***Purification of human and mouse neutrophils.*** All mice were bred and maintained under pathogen-free conditions at an American Association for the Accreditation of Laboratory Animal Care accredited animal facility at the NIAID and housed in accordance with the procedures outlined in the Guide for the Care and Use of Laboratory Animals under an animal study proposal approved by the NIAID Animal Care and Use Committee. Mouse neutrophils were purified from bone marrow as previously published (Wen et al., 2018). Briefly, one end of the hind legs of C57/B6 mice was cut open and bone marrow was collected by centrifugation at 8600 × *g* for 10 s. Red blood cells were removed with ACK buffer (Lonza) at room temperature. Cells were counted and subjected to a neutrophil isolation procedure using a mouse neutrophil isolation kit (Miltenyi Biotec, order No. 130097658). Collected neutrophils were stored in PBS supplemented with 2% FBS and 2 mM EDTA at 4°C. The purity of neutrophils was determined by staining the cells with mouse Gr-1/Ly-6G PE-conjugated antibody and FITC-conjugated CD11b antibody (R&D systems). The purity was above 95%.

Peripheral blood samples from healthy U.S. adults were obtained from the National Institutes of Health (NIH) Department of Transfusion Medicine under an NIH Institutional Review Board-approved protocol (99-CC-0168) with informed consent. Human neutrophils were purified from human blood as previously reported (Xu et al., 2015a). Whole blood (150 ml) was drawn from healthy volunteers at the Blood Bank of the National Institutes of Health. Coagulation was prevented by heparin. The majority of the red blood cells were removed by dextran (0.2 g/l; GE Healthcare, Pittsburgh, PA) sedimentation (30 min, room temperature). Upper phases containing white blood cells were collected and washed twice in phosphate-buffered saline (PBS). Cells were resuspended in PBS and layered on top of a five-step Percoll gradient (65, 70, 75, 80, and 85%; Sigma-Aldrich) in 15-ml conical tubes. After centrifugation ($800 \times g$, 20 min, room temperature), the 70/75/80% Percoll layers containing granulocytes were collected and washed twice in PBS. The collected neutrophils were resuspended in PBS, the cell concentration was determined, and the cells were kept at room temperature until use. Cell viability was determined by trypan blue dye extrusion, and the results were >98% viable neutrophils. The purity of the preparations was determined by Wright-Giemsa staining, and the yield was >95% neutrophil granulocytes.

**Reagents and antibodies**

$\alpha$-PKC$\alpha$, -p-PKC$\alpha$(S657/Y658), -p-PKC(pan)( $\beta$IIS660), -p-PKC(pan)( $\zeta$T410), -p-PKC$\alpha$/$\beta$II(T638/T641), -CD11, and -FPR1 antibodies from Cell Signaling Technology (Beverly, MA); $\alpha$-PLC$\beta$2 antibodies from LifeSpan BioScience, Inc. (Seattle, WA); $\alpha$-actin and -PLC$\gamma$1 antibodies from Thermo Scientific (Barrington, IL); $\alpha$-PLC$\beta$2 and -PLC$\beta$3 antibodies from Abcam (Cambridge, MA); $\alpha$-PLC$\varepsilon$1 antibody from Biocompare (San Francisco, CA); HRP-conjugated anti-mouse or anti-rabbit IgG from Jackson ImmunoResearch (West Grove, PA); and all tissue culture reagents from Invitrogen (Carlsbad, CA).

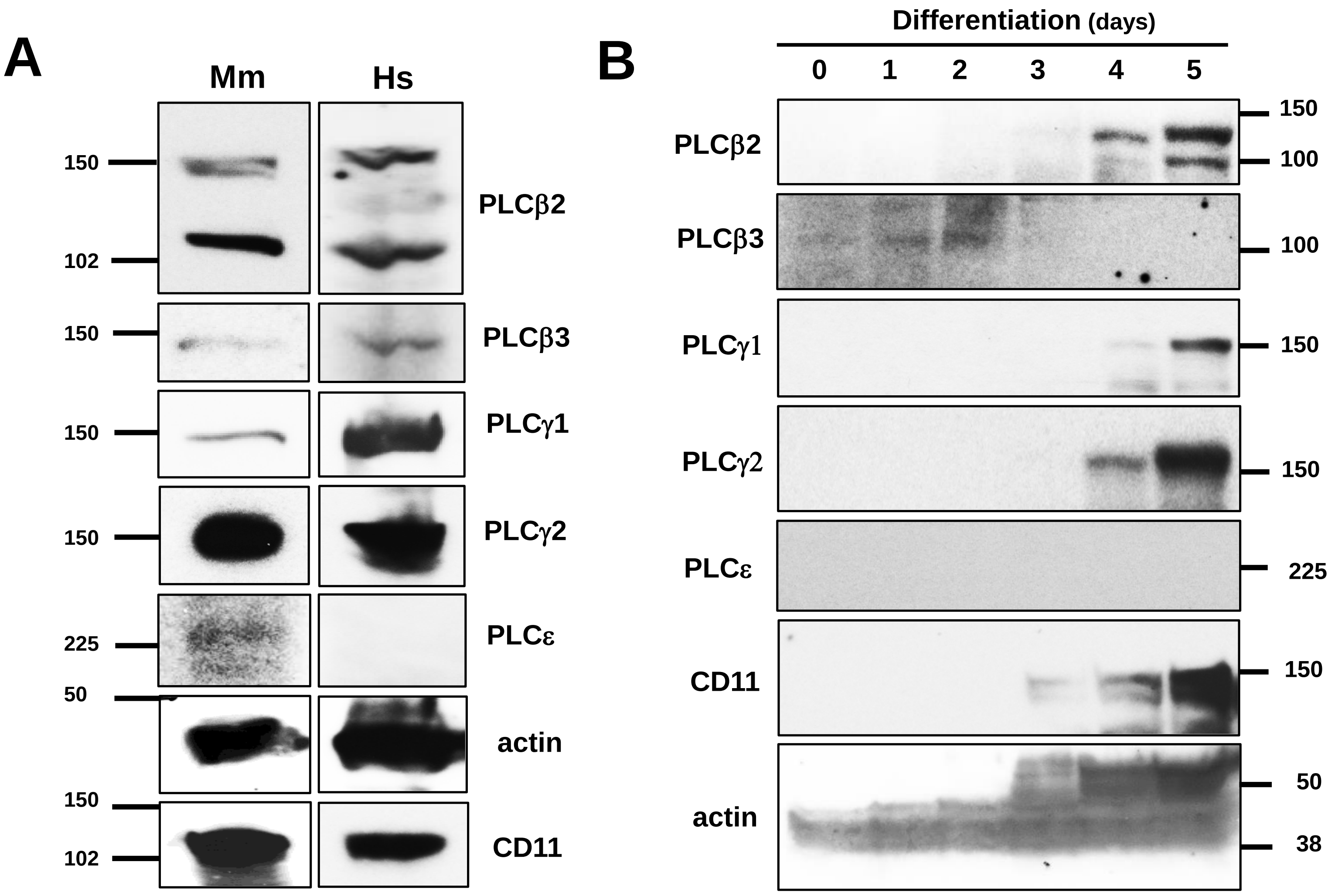
A
Mm
Hs
150
102
PLCβ2
150
PLCβ3
150
PLCγ1
150
PLCγ2
225
PLCε
50
actin
150
102
CD11
B
Differentiation (days)
0
1
2
3
4
5
PLCβ2
150
100
PLCβ3
100
PLCγ1
150
PLCγ2
150
PLCε
225
CD11
150
actin
50
38

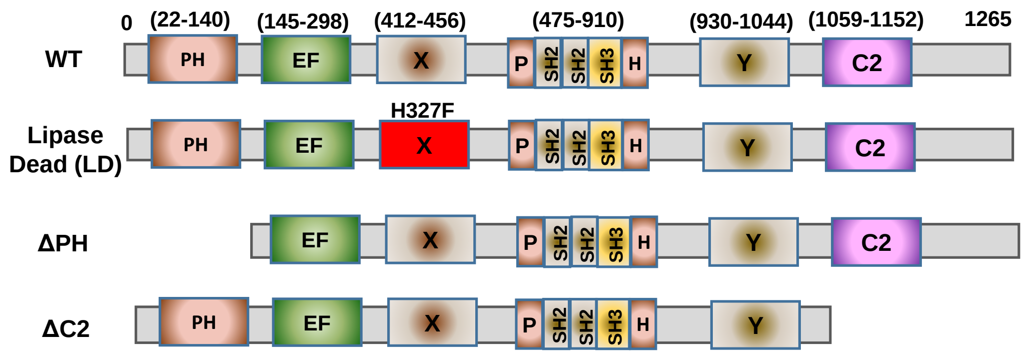
0
(22-140)
(145-298)
(412-456)
(475-910)
(930-1044)
(1059-1152)
1265
WT
PH
EF
X
P
SH2
SH2
SH3
H
Y
C2
H327F
Lipase
Dead (LD)
ΔPH
ΔC2

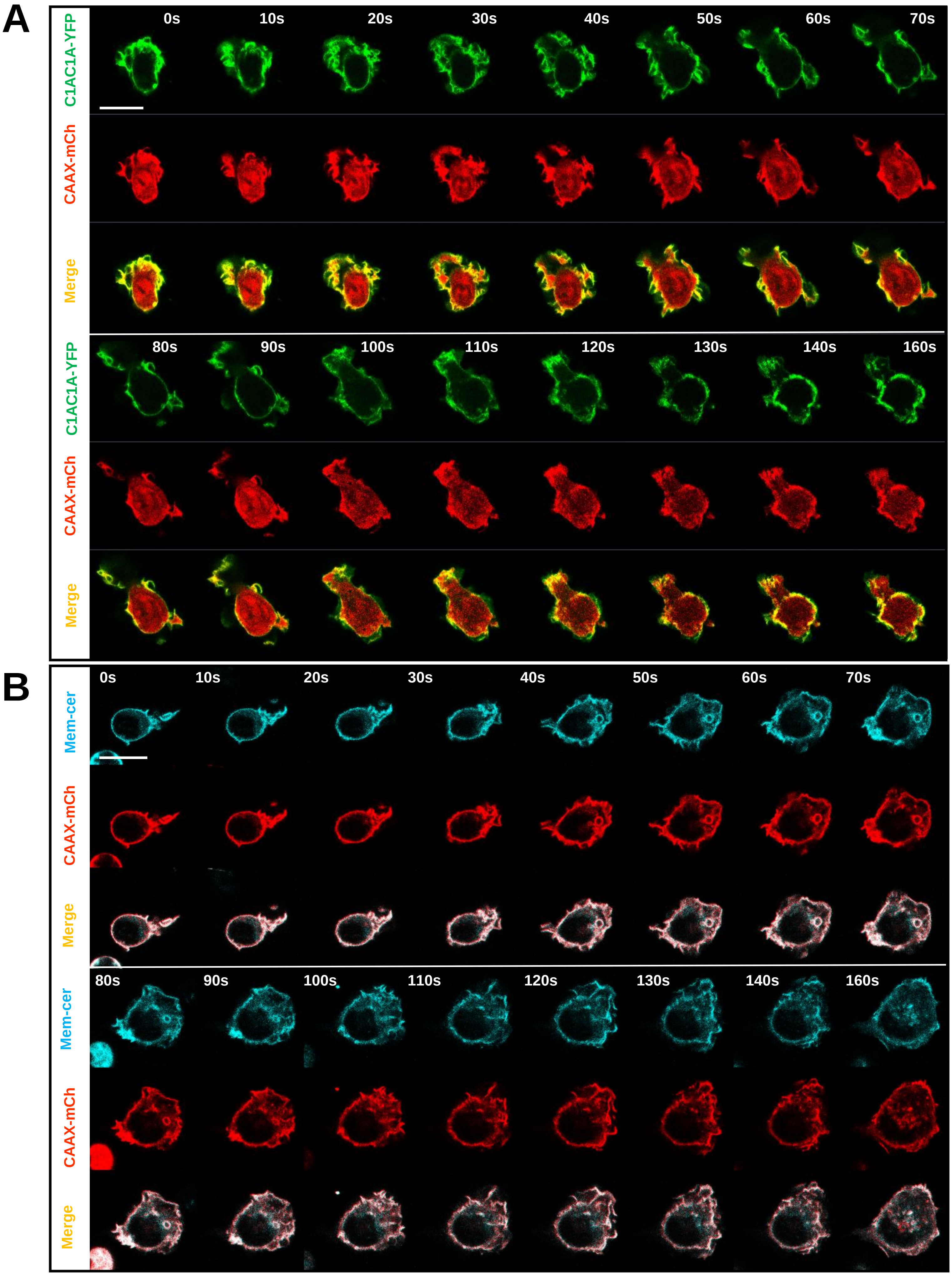
A
C1AC1A-YFP
CAAX-mCh
Merge
0s
10s
20s
30s
40s
50s
60s
70s
80s
90s
100s
110s
120s
130s
140s
160s
B
Mem-cer
CAAX-mCh
Merge

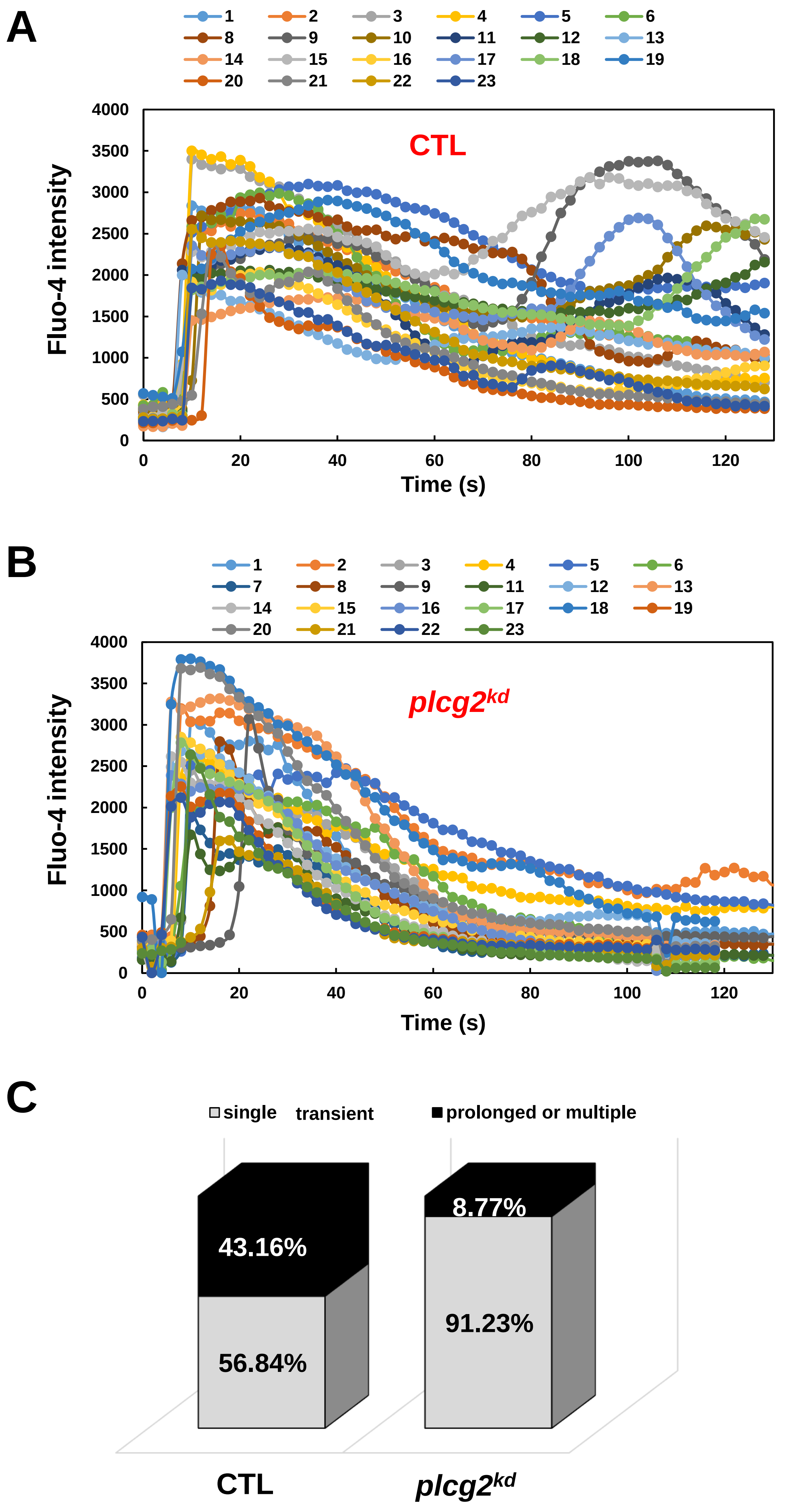
A
1
2
3
4
5
6
CTL
Fluo-4 intensity
Time (s)
B
1
2
3
4
5
6
7
8
9
plcg2kd
Fluo-4 intensity
Time (s)
C
single transient
prolonged or multiple
43.16%
56.84%
8.77%
91.23%
CTL
plcg2kd

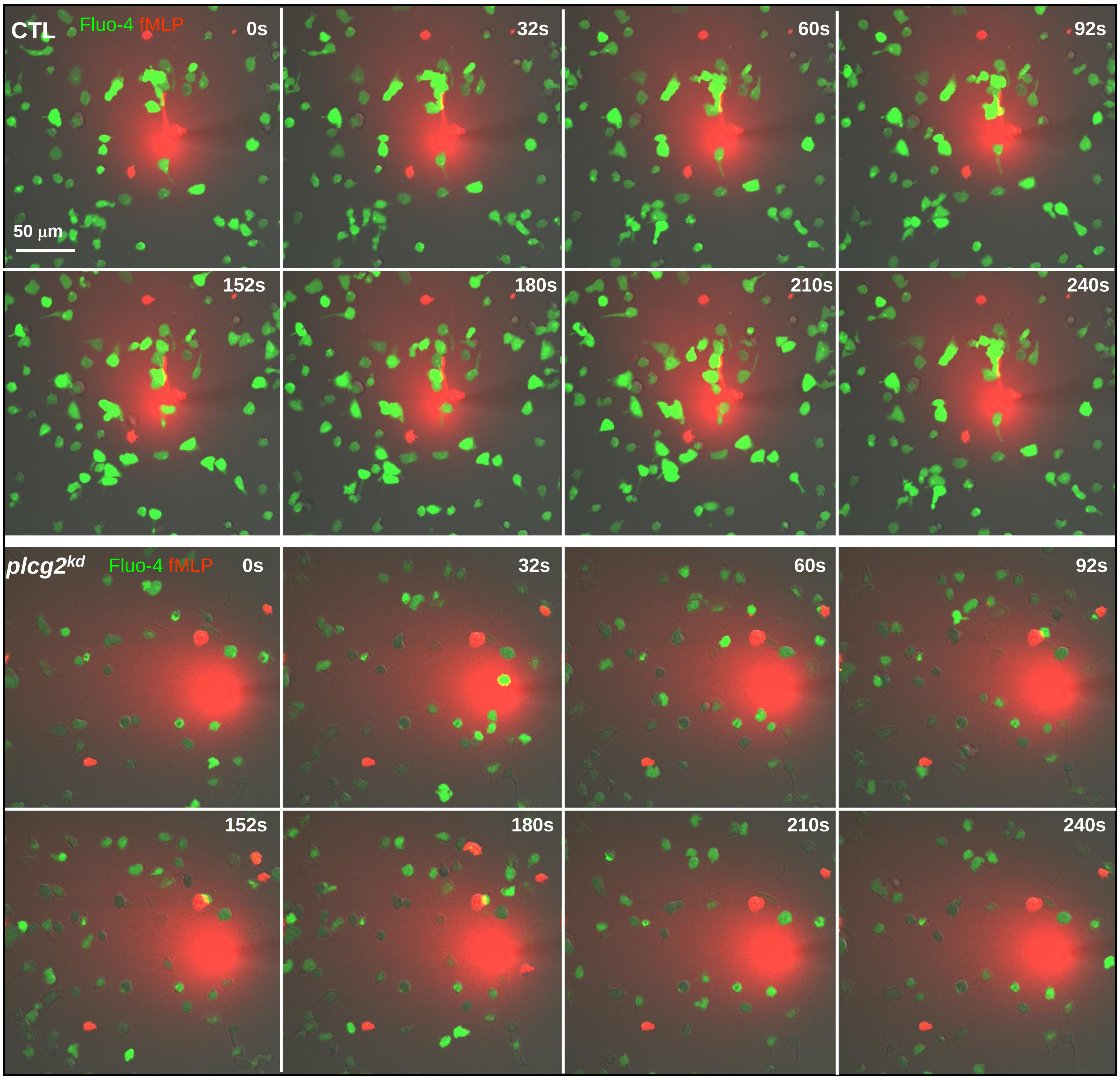
CTL
Fluo-4 fMLP
0s
32s
60s
92s
50 μm
152s
180s
210s
240s
plcg2kd
Fluo-4 fMLP
0s
32s
60s
92s
152s
180s
210s
240s

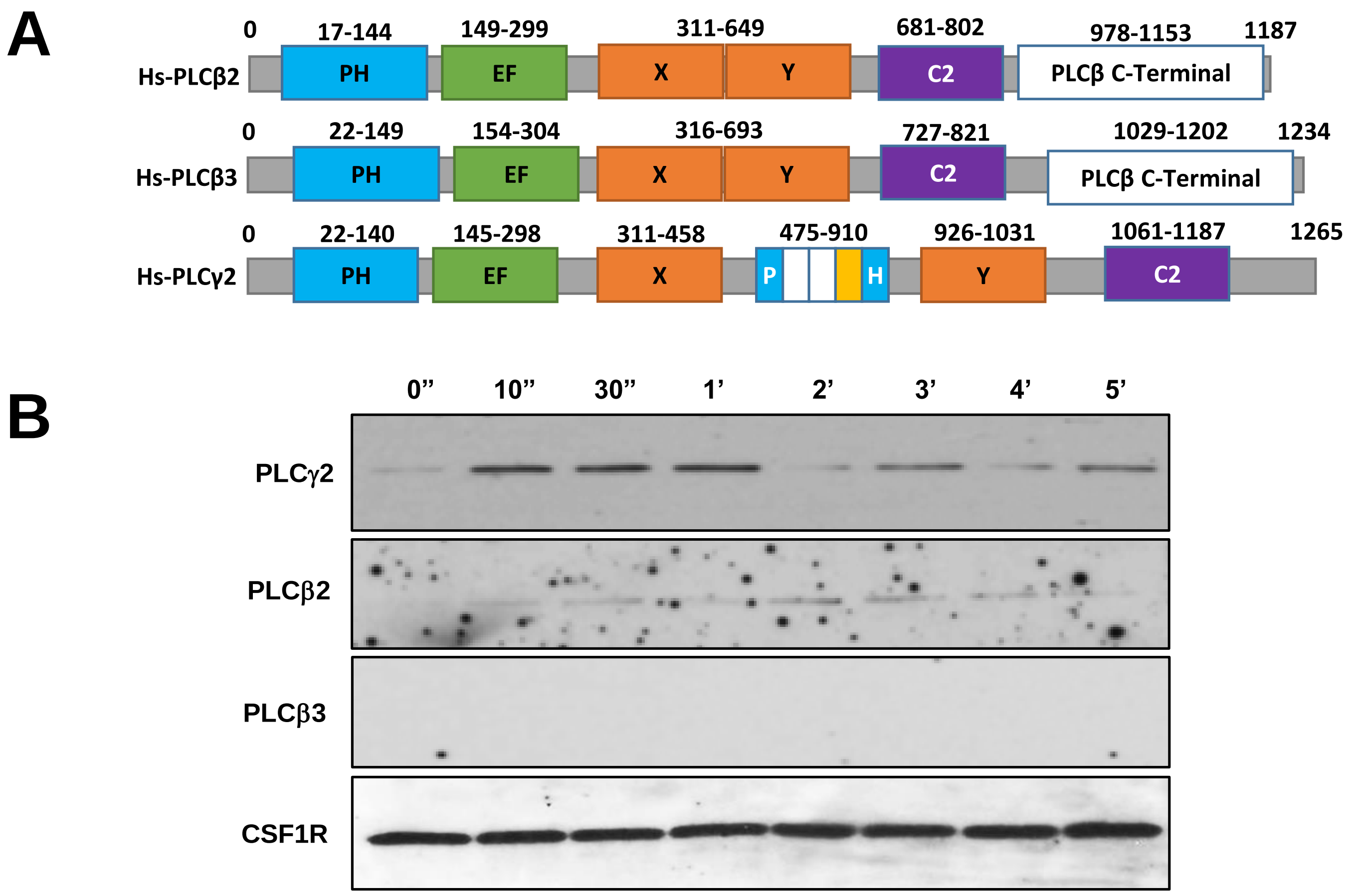

C

Conserved PIP-binding Residues　Conserved Hydrophobic Residues

PIP3 Binding

```
PH-HsAKT      4-VAIVKEGWLHKRGE-YIK----TWRPRYFLLKNDGTFIGYKER-41
PH-HsBTK      40----LESIFLKRSQQKKKTSPLNFKKRLFLL--TVHKLSYYEY-76
PH-HsGRP1     265-PDREGWLLKLGGGRVK----TWKRRWFIL--TDNCLYYFEY-299
PH-HsPLCγ2    22----LELGTVMTVFSFRKST---PERRTVQVIMETRQVAWSKT-57
PH-HsPLCβ2    17----LSQGERFIKWDDETTV---ASPVILRVDPKGYYLYWTYQ-53
PH-HsPLCβ3    22---LRRGSKFIKWDEETSS----RNLVTLRVDPNGFFLYWTGP-57
```

PIP2 BindingC

```
PH-HsPLCδ1    43---LLKGSQLLKVK-SSSW-----RRERFYKLQEDCKTI-WQES-76
PH-HsPLCγ1    115--MQAGCELKKVR-PNS-----RIYNRFFTLDTDLQALRWEP-148
PH-HsPLCγ2    22---LELGTVMTVFSFRKST----PERRTVQVIMETRQVAWSKT-57
PH-HsPLCβ2    17---LSQGERFIKWDDETTV-----ASPVILRVDPKGYYLYWTYQ-53
PH-HsPLCβ3    22---LRRGSKFIKWDEETSS-----RNLVTLRVDPNGFFLYWTGP-57
```

D

$Ca^{2+}$ Binding

```
C2-HsSYT1     304---DVGGLSDPYVKIHLMQNGKRLKKKKTTIKKNTLNP[16]KVQVVVTVLDYDKIGKND-372
C2-HsPKCβ1    187---DPNGLSDPYVKLKLIPDPKSESKQKTKTIKCSLNP[15]DRRLSVEIWDWDLTSRND-254
C2-HsPLCγ2    1075-KLGRSIADPFVEVEICGAEYDNNKFKTTVVNDNGLS[18]LAFLRFLVYEEDMFS--DPN-1146
C2-HsPLCβ2    696---ERSVRTYVEVDMFGLPGDPKRRYRTKLSPSTNS[20]-ASLRVAVMEEG----NK---761
C2-HsPLCβ3    740---DRKVGIYVEVDMFGLPVDTRRKYRTRTSQGNSF[20]-ASLRIAAFEEG----GK-804
```

**A**

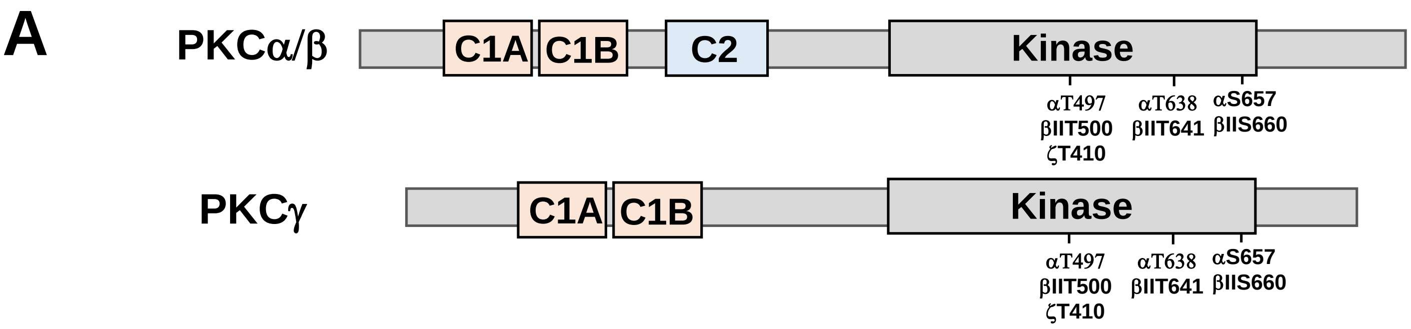

**B**

```
        VPVPXXDX-------EANXELRQKFERAKIXXXGXKVPSPSXXXTXXK---FXXSXX-DRLKLTDFNFLMVLGKGSFGKV
                330       340       350       360       370       380       390       400
PKCγII  VPVADADNCSLLQKFEACNYPLELYERVRMGPSSSPIPSPSPSPTDPKRCFFGASP--GRLHISDFSFLMVLGKGSFGKV  365
PKCα    VPIPEGDE-------EGNMELRQKFEKAKLGPAGNKVISPSEDRK-------QPSNNLDRVKLTDFNFLMVLGKGSFGKV  353
PKCβII  VPVPPEGS-------EANEELRQKFERAKISQ-GTKVPEEKTTNTVSK---FDNNGNRDRMKLTDFNFLMVLGKGSFGKV  356
PKCβI   VPVPPEGS-------EANEELRQKFERAKISQ-GTKVPEEKTTNTVSK---FDNNGNRDRMKLTDFNFLMVLGKGSFGKV  356
PKCζI   -------------------------MDGIKISQG--------------------------LGLQDFDLIRVIGRGSYAKV  266
PKCγI   VPVADADNCSLLQKFEACNYPLELYERVRMGPSSSPIPSPSPSPTDPKRCFFGASP--GRLHISDFSFLMVLGKGSFGKV  365

        MLAERKGTDELYAIKILKKDVVIQDDDVXCTMVEKRVLAL----XXGKPPFLTQLHSCFQTXDRLYFVMEYVNGGDLMYH
                410       420       430       440       450       460       470       480
PKCγ2   MLAERRGSDELYAIKILKKDVIVQDDDVDCTLVEKRVLALGGRGPGGRPHFLTQLHSTFQTPDRLYFVMEYVTGGDLMYH  445
PKCα    MLADRKGTEELYAIKILKKDVVIQDDDVECTMVEKRVLAL-----LDKPPFLTQLHSCFQTVDRLYFVMEYVNGGDLMYH  428
PKCβ2   MLSERKGTDELYAVKILKKDVVIQDDDVECTMVEKRVLAL-----PGKPPFLTQLHSCFQTMDRLYFVMEYVNGGDLMYH  431
PKCβ1   MLSERKGTDELYAVKILKKDVVIQDDDVECTMVEKRVLAL-----PGKPPFLTQLHSCFQTMDRLYFVMEYVNGGDLMYH  431
PKCζ1   LLVRLKKNDQIYAMKVVKKELVHDDEDIDWVQTEKHVFEQASSNP-----FLVGLHSCFQTTSRLFLVIEYVNGGDLMFH  341
PKCγ1   MLAERRGSDELYAIKILKKDVIVQDDDVDCTLVEKRVLALGGRGPGGRPHFLTQLHSTFQTPDRLYFVMEYVTGGDLMYH  445

        IQQVGKFKEPHAVFYAAEIAIGLFFLHXXGIIYRDLKLDNVMLDXEGHIKIXDFGMCKENXXXGVTTRTFCGTPDYIAPE
                490       500       510       520       530       540       550       560
PKCγ2   IQQLGKFKEPHAAFYAAEIAIGLFFLHNQGIIYRDLKLDNVMLDAEGHIKITDFGMCKENVFPGTTTRTFCGTPDYIAPE  525
PKCα    IQQVGKFKEPQAVFYAAEISIGLFFLHKRGIIYRDLKLDNVMLDSEGHIKIADFGMCKEHMMDGVTTRTFCGTPDYIAPE  508
PKCβ2   IQQVGRFKEPHAVFYAAEIAIGLFFLQSKGIIYRDLKLDNVMLDSEGHIKIADFGMCKENIWDGVTTKTFCGTPDYIAPE  511
PKCβ1   IQQVGRFKEPHAVFYAAEIAIGLFFLQSKGIIYRDLKLDNVMLDSEGHIKIADFGMCKENIWDGVTTKTFCGTPDYIAPE  511
PKCζ1   MQRQRKLPEEHARFYAAEICIALNFLHERGIIYRDLKLDNVLLDADGHIKLTDYGMCKEGLGPGDTTSTFCGTPNYIAPE  421
PKCγ1   IQQLGKFKEPHAAFYAAEIAIGLFFLHNQGIIYRDLKLDNVMLDAEGHIKITDFGMCKENVFPGTTTRTFCGTPDYIAPE  525

        IIAYQPYGKSVDWWAFGVLLYEMLAGQPPFDG-------EDEDELFQSIMEHNVXYPKSLSKEAVAICKGXXTKHPGKRL
                570       580       590       600       610       620       630       640
PKCγ2   IIAYQPYGKSVDWWSFGVLLYEMLAGQPPFDG-------EDEEELFQAIMEQTVTYPKSLSREAVAICKGFLTKHPGKRL  598
PKCα    IIAYQPYGKSVDWWAYGVLLYEMLAGQPPFDG-------EDEDELFQSIMEHNVSYPKSLSKEAVSICKGLMTKHPAKRL  581
PKCβ2   IIAYQPYGKSVDWWAFGVLLYEMLAGQAPFEG-------EDEDELFQSIMEHNVAYPKSMSKEAVAICKGLMTKHPGKRL  584
PKCβ1   IIAYQPYGKSVDWWAFGVLLYEMLAGQAPFEG-------EDEDELFQSIMEHNVAYPKSMSKEAVAICKGLMTKHPGKRL  584
PKCζ1   ILRGEEYGFSVDWWALGVLMFEMMAGRSPFDIITDNPDMNTEDYLFQVILEKPIRIPRFLSVKASHVLKGFLNKDPKERL  501
PKCγ1   IIAYQPYGKSVDWWSFGVLLYEMLAGQPPFDG-------EDEEELFQAIMEQTVTYPKSLSREAVAICKGFLTKHPGKRL  598

        GCGPEGE-RDIXEHAFFRXIDWEKLERKEIQPPFKPKXCG-RXXENFDKFFTRAPPXLTPPDXLVIANIDQSXFEGFSYV
                650       660       670       680       690       700       710       720
PKCγ2   GSGPDGE-PTIRAHGFFRWIDWERLERLEIPPPFRPRPCG-RSGENFDKFFTRAAPALTPPDRLVLASIDQADFQGFTYV  676
PKCα    GCGPEGE-RDVREHAFFRRIDWEKLENREIQPPFKPKVCGK-GAENFDKFFTRGQPVLTPPDQLVIANIDQSDFEGFSYV  659
PKCβ2   GCGPEGE-RDIKEHAFFRYIDWEKLERKEIQPPYKPKACG-RNAENFDRFFTRHPPVLTPPDQEVIRNIDQSEFEGFSFV  662
PKCβ1   GCGPEGE-RDIKEHAFFRYIDWEKLERKEIQPPYKPKARDKRDTSNFDKEFTRQPVELTPTDKLFIMNLDQNEFAGFSYT  663
PKCζ1   GCRPQTGFSDIKSHAFFRSIDWDLLEKKQALPPFQPQITDDYGLDNFDTQFTSEPVQLTPDDEDAIKRIDQSEFEGFEYI  581
PKCγ1   GSGPDGE-PTIRAHGFFRWIDWERLERLEIPPPFRPRPCG-RSGENFDKFFTRAAPALTPPDRLVLASIDQADFQGFTYV  676

        NPXFVHPXARX-X--------------------
                730       740       750
PKCγ2   NPDFVHPDARSPTSPVPVPVM                                                              697
PKCα    NPQFVHPILQSAV                                                                      672
PKCβ2   NSEFLKPEVK--S                                                                      673
PKCβ1   NPEFV---IN--V                                                                      671
PKCζ1   NPLLLSTEE-----------------------SV                                                 592
PKCγ1   NPDFVHPDARSPTSPVPVPVISCTPAFQLCPRGF                                                 710
```

Xu et al., Figure S8

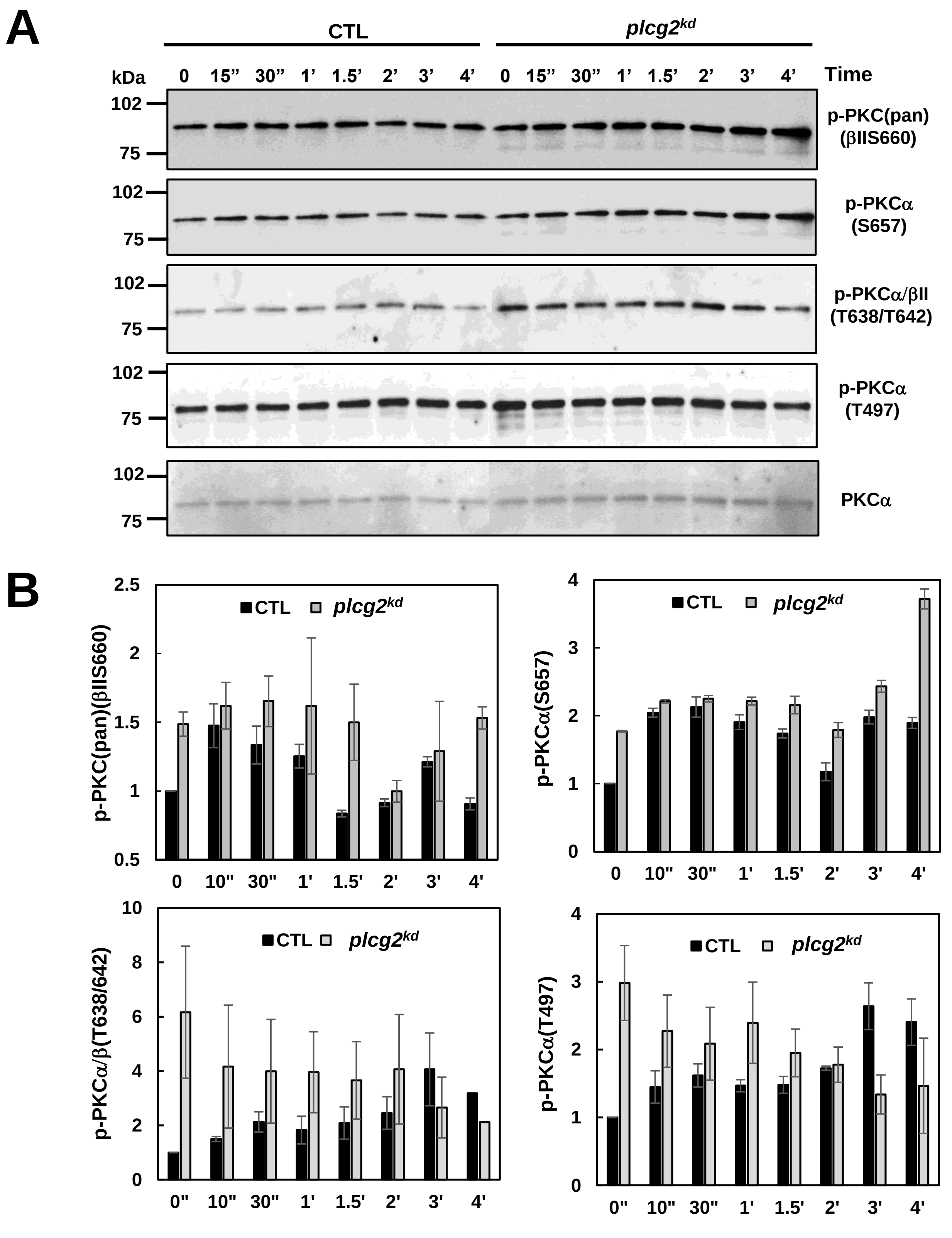
